# *Maleness-on-the-Y* (*MoY*) orchestrates male sex determination in major agricultural fruit fly pests

**DOI:** 10.1101/533646

**Authors:** Angela Meccariello, Marco Salvemini, Pasquale Primo, Brantley Hall, Panagiota Koskinioti, Martina Dalíková, Andrea Gravina, Michela Anna Gucciardino, Federica Forlenza, Maria-Eleni Gregoriou, Domenica Ippolito, Simona Maria Monti, Valeria Petrella, Maryanna Martina Perrotta, Stephan Schmeing, Alessia Ruggiero, Francesca Scolari, Ennio Giordano, Konstantina T. Tsoumani, Frantisek Marec, Nikolai Windbichler, Javaregowda Nagaraju, Kallare P. Arunkumar, Kostas Bourtzis, Kostas D. Mathiopoulos, Jiannis Ragoussis, Luigi Vitagliano, Zhijian Tu, Philippos Aris Papathanos, Mark D. Robinson, Giuseppe Saccone

**Author notes:** These authors contributed equally to this work (Angela Meccariello, Marco Salvemini).

## Abstract

In insects, rapidly evolving primary sex-determining signals are transduced by a conserved regulatory module producing sex-specific proteins that direct sex determination and sexual differentiation^1-4^. In the agricultural pest *Ceratitis capitata* (medfly), a Y-linked maleness factor (*M*) is thought to repress the autoregulatory splicing of *transformer* (*Cctra*), which is required in XX individuals to establish and maintain female sex determination^5,6^. Despite previous attempts of isolating Y-linked genes using the medfly whole genome, the *M* factor has remained elusive^7^. Here, we report the identification of a Y-linked gene, ***M****aleness-****o****n the-****Y*** (*MoY*), and show that it encodes a small novel protein which is both necessary and sufficient for medfly male sex determination. Transient silencing of *MoY* in XY individuals leads to the development of fertile females while transient expression of *MoY* in XX individuals results in fertile males. Notably, a cross between these sex reverted individuals gives rise to both fertile males and females indicating that a functional *MoY* can be maternally transmitted. In contrast to the diversity of *M* factors found in dipteran species^8-11^, we discovered *MoY* orthologues in seven other Tephritid species spanning ∼111 millions of years of evolution (Mya). We confirmed their male determining function in the olive fly (*Bactrocera oleae*) and the oriental fruit fly (*Bactrocera dorsalis*). This unexpected conservation of the primary *MoY* signal in a large number of important agricultural pests^12^ will facilitate the development of transferable genetic control strategies in these species, for example sterile male releases or sex-ratio-distorting gene drives.

## Introduction

The Tephritidae family includes 5000 species, dozens of which are invasive and highly relevant agricultural pests of fruits and crop. *Ceratitis capitata* (medfly) is one of the most destructive members of this taxon, affecting over 200 plant species. The most successful method to control the medfly, the Sterile Insect Technique (SIT)^13^, involves the continuous mass-release of laboratory-reared sterilized males that suppress, or even eradicate wild populations by mating with wild females. A key determinant to the success of SIT programs, has been the close physical linkage of selectable traits to the Y-linked *M* factor in so called genetic sexing strains, that were generated to enable sex separation on a massive scale and thus the mass male release^14^. The development of similar strains in other Tephritid pest species however has been difficult, inhibiting the transfer of SIT or related genetic strategies. Therefore identifying the *M* factor of the medfly and documenting its evolutionary origin and conservation in related pests also holds significant promise for establishing genetic control strategies in species where efficient population control methods are currently needed.

## Results

We conducted the medfly *M* factor search by: 1) focusing on the 4-8 hours after egg laying (AEL) time window to assess embryonic transcription, the period when male sex determination is first established^15^; 2) producing XX-only embryonic transcriptomes used as a control in the search for Y-specific sequences; 3) developing a PacBio-based male genome assembly from the *Fam18* medfly strain^6^, that has a short Y chromosome; and 4) searching for conservation of putative *M* factors in another Tephritidae, the olive fly *Bactrocera oleae*. We found 19 candidate Y-chromosome derived transcripts that are expressed specifically in mixed-sex embryos, with no assigned reads from XX-only embryos (Fig. 1a; Extended Data Table 1; Supplementary Note 1). Seven of these transcripts did not map to the *Fam18* male genome assembly and were excluded. Sequence similarity searches by BLASTn showed that three out of 12 transcripts had hits to XY but not XX embryonic transcripts from the related *B. oleae.* Furthermore, one of these (DN40292_c0_g3_i1) corresponds to one of the 28 transcripts previously identified by CQ approach^16^ in a preliminary screen (Supplementary Note 2). This transcript mapped on a predicted Y-linked scaffold in the *Fam18* genome assembly (Supplementary Note 3 and 4) and lacked paralogs elsewhere in the genome, suggesting that it may be single-copy on the Y. Functional analysis (below) confirmed that the corresponding gene is the medfly *M* factor and was thus named ***M****aleness-****o****n-the-****Y*** (*MoY*). *MoY* is a short intronless gene located on the long arm of the Y chromosome in proximity to the centromere (Fig. 1b) in a region that contains 9 other transcriptional units, most of which are repetitive and largely overlapping on two opposite strands (Fig. 1c; Supplementary Notes 1, 3 and 4). *MoY* expression begins at 2-3h AEL, prior to embryonic cellularization, with a transient peak at 15h and becomes undetectable by 48h (Fig. 1d).

**Figure 1.**
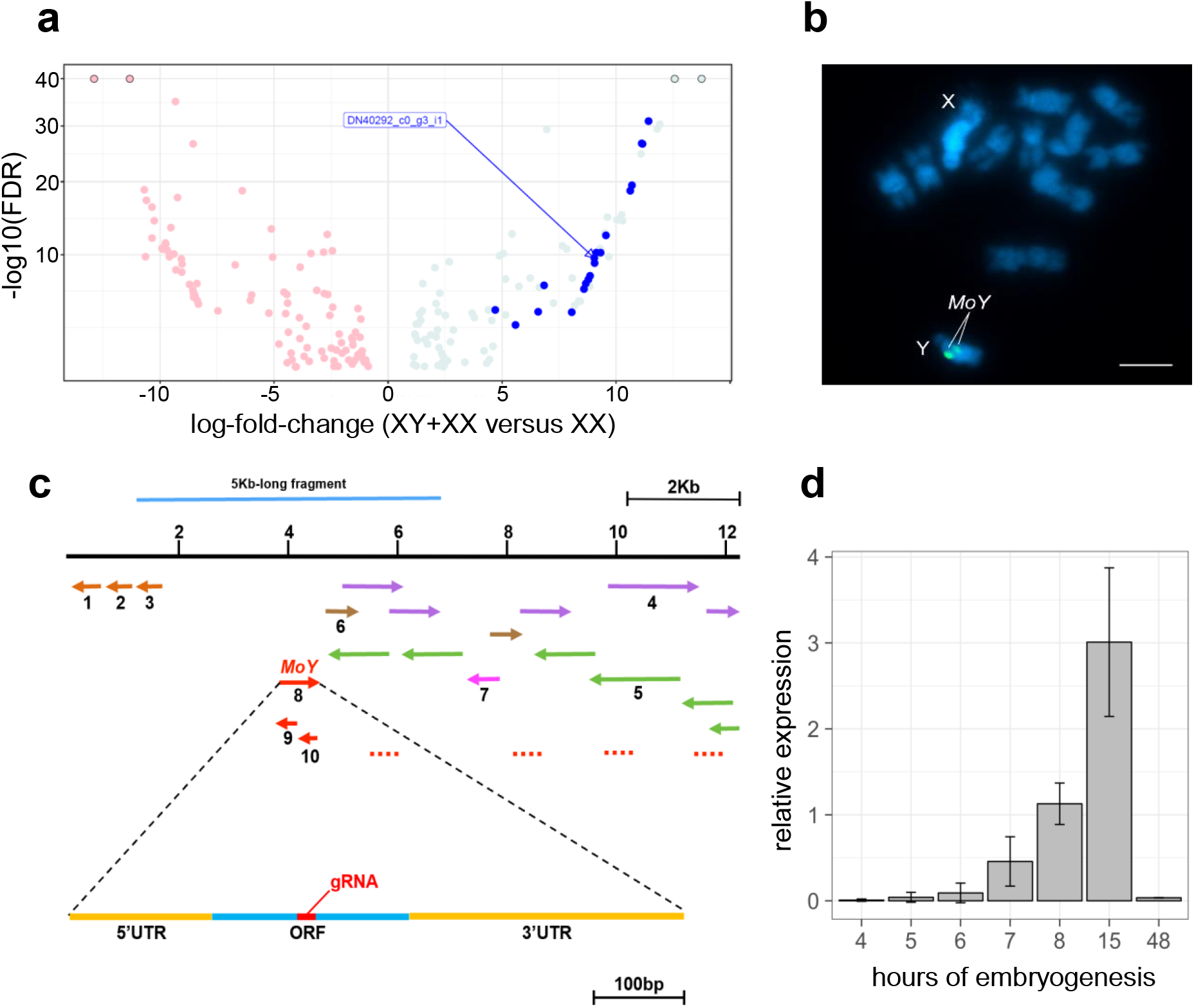
Identification of the medfly Y-linked Male determining factor (*MoY*), a single copy gene active during early embryogenesis. **a**, Volcano plot of the 195 differentially expressed transcripts at 4-8 h AEL, with FDR < 0.05. Pink dots indicate female-biased transcripts while azure dots indicate male-biased transcripts. Blue dots indicated transcripts with a CQ value of 0 (putative Y-linked transcripts). The transcript corresponding to *MoY* (DN40292_c0_g3_i1) is indicated by arrow. Dots at the top with a black border indicate transcripts where the FDR values were lower, but were set to 1e-40 for visualisation. **b**, Fluorescence *in situ* hybridization of *MoY* (2 green signals) on mitotic chromosomes stained with DAPI (blue); twin signals (because of two sister chromatids) locate *MoY* on the long arm of the Y chromosome near the centromere. **c**, 12 Kb long genomic region (from PacBio Canu assembly; GenBank accession number: MK330842) showing in red the *MoY* transcript (8) antisense RNA (in red 9-10) and weakly related sequences (in red discontinuous lines), in violet (4) and green (5) a repeated sequence transcribed in both strands, in light brown 3 unrelated transcripts (1-3), in brown (6) and light violet (7) other transcriptional novel units. **d**, *E*xpression profile of *MoY* during medfly embryogenesis. A peak at 15 h is observed.

Transient silencing of *MoY* by embryonic RNAi microinjections resulted in the loss of male-specific *Cctra* transcripts in 8h AEL embryos (Extended Data Table 2; Supplementary Table 1). When we assessed *Cctra* splicing in injected 3-day-old XY larvae and adults we found a switching to the female-specific autoregulatory pattern^5^(Fig. 2a-b; Extended Data Fig. 1a-b;). Among adults, 38% of the XY individuals (14/37) displayed complete phenotypic feminization and 19% (7/37) had intersex phenotypes (Fig. 2c). Phenotypic assignment of sex was based on the presence/absence of the pair of male-specific spatulae and of the female-specific ovipositor. Individuals lacking the female-specific ovipositor and male-specific spatulae, or having both, were phenotypically classified as intersexes. To evaluate the fertility of these feminized XY individuals, crosses were established to XX fertile males, which were generated by silencing of *Cctra*^5^. This crossing strategy simplified the tracking of any Y chromosomes that were transmitted maternally (Extended Data Fig. 3). Among the 13 recovered XY females, 2 were fertile and transmitted their Y chromosome to male progeny (Supplementary Table 2, female n. 26, Extended Data Fig. 3, Extended Data Table 2). To our knowledge, this is the first experiment in an animal species with heteromorphic XY chromosomes, in which a functional Y chromosome was transmitted maternally (Extended Data Figure 3a).

**Figure 2.**
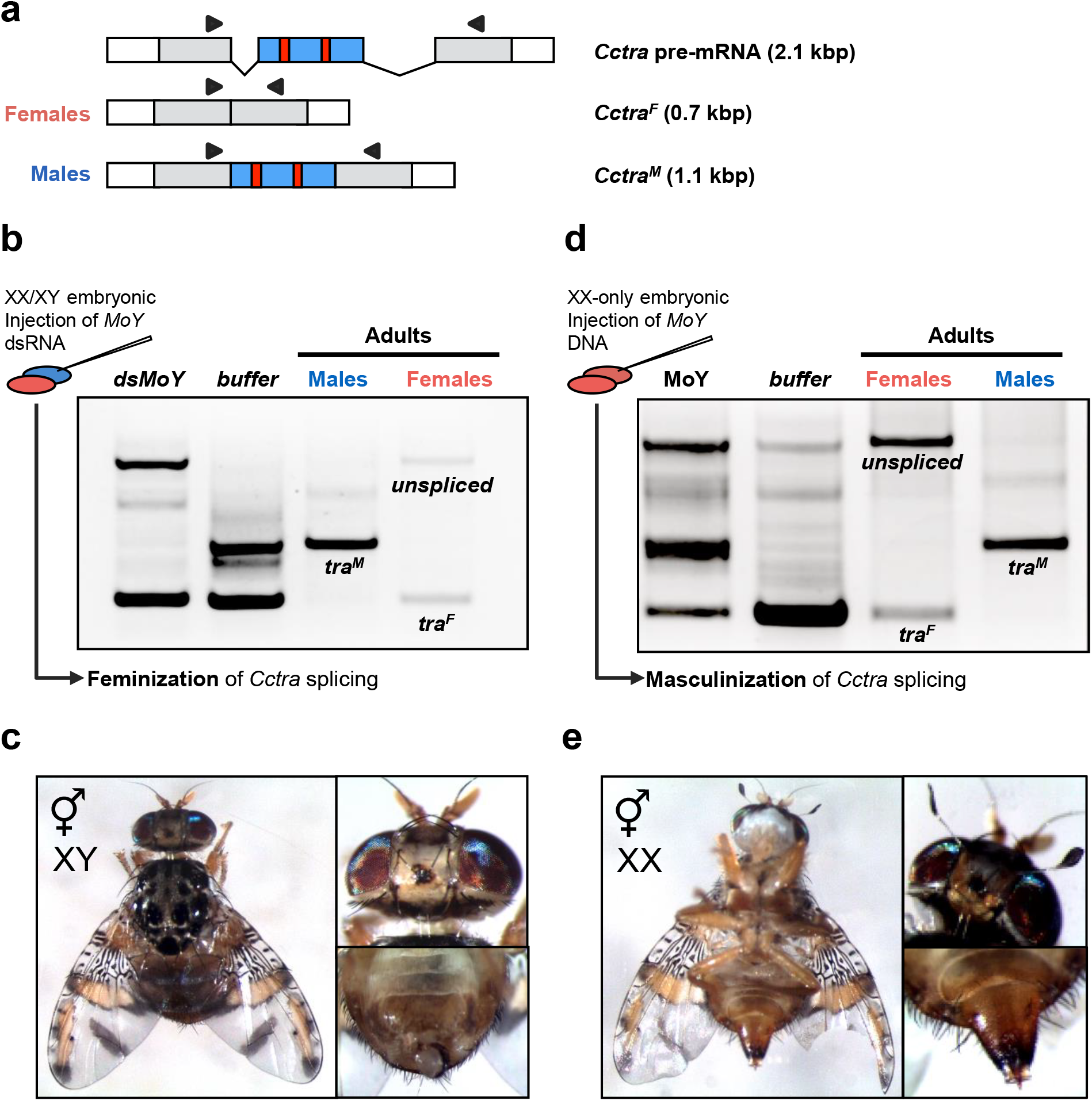
*MoY* is necessary and sufficient for male sex determination. **a**, Scheme of *Cctra* gene and male- and female-specific transcripts. Coding region is in gray, male-specific exons are in blue, red bars indicate stop codons in the male-specific exon. Black arrows indicate primers used for RT-PCR. **b**, Transient embryonic *MoY* RNAi is sufficient to feminize XY individuals. RT-PCR analysis showing splicing patterns of *Cctra* in 8 h old embryos, following dsRNA *MoY* or buffer injections at 0-1 h AEL, compared to wild type adult flies. **c**, An example of an XY adult intersex developed from *MoY* dsRNA injected embryos (Table 1, 1#) showing male genitalia and a female-like head, lacking the orbital setae (for wild type reference see Supplementary Fig. 2). **d**, XX-only embryonic injections of genomic DNA containing *MoY* is sufficient to induce male-specific splicing of *Cctra,* but not the buffer injection, compared to wild type adult flies. **e,** An example of a XX adult intersex showing female ovipositor and a male-like head, with orbital setae.

**Figure 3.**
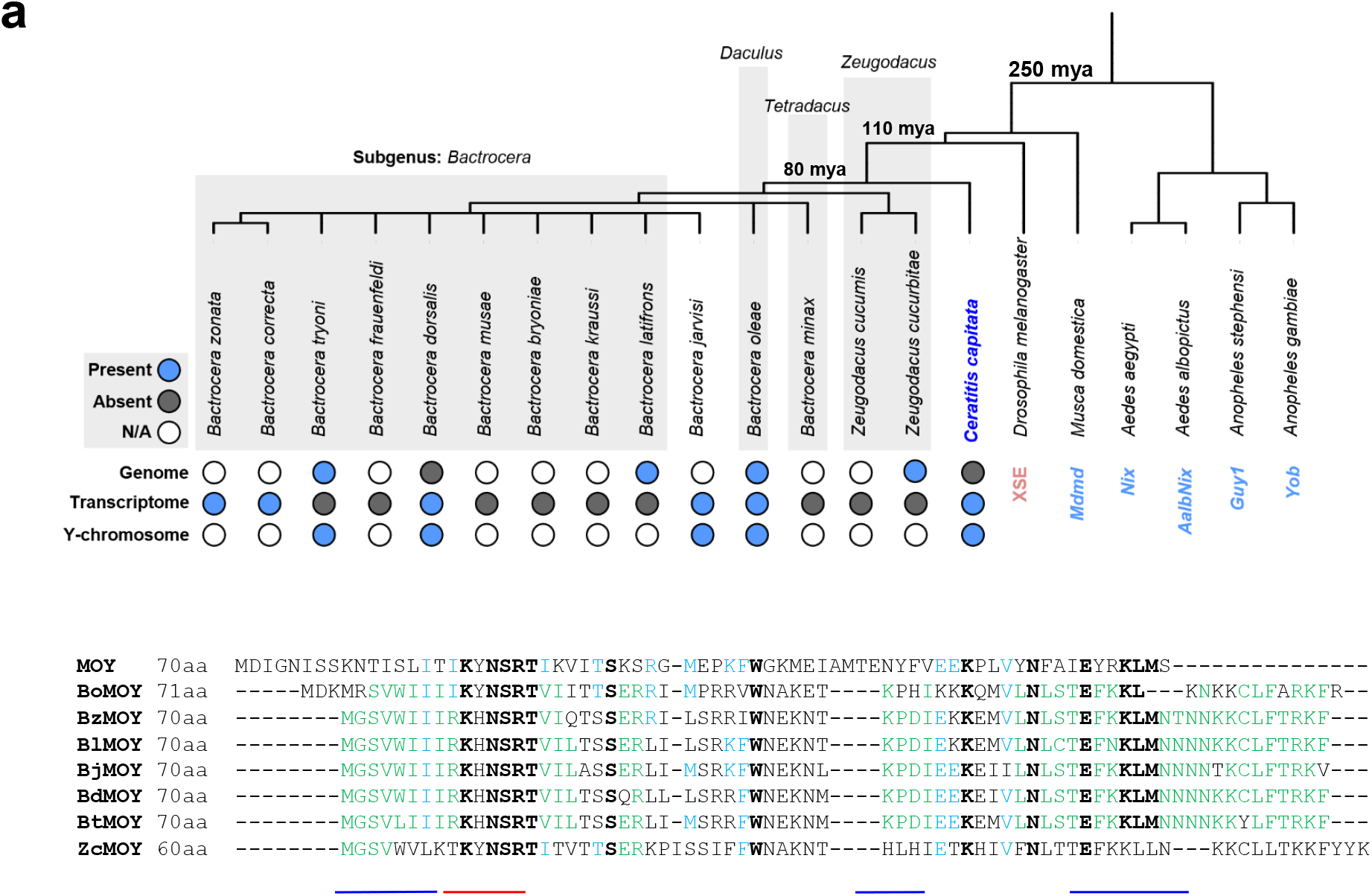
*MoY* gene is structurally and functionally conserved in Tephritidae species. **a**, Phylogenetic tree of dipteran species18, in which male determining genes or XSE (*Drosophila*) primary sex determining signals have been molecularly characterized. *MoY* orthologues were searched in 14 Tephritidae species, and found in 8 either in genome and/or transcriptome available databases at NCBI (Supplementary Information Notes 5-12) (The lack of hits in 6 out of 14 assembled transcriptomes is likely due to insufficient sequencing coverage). **b**, Manual alignment of 8 MOY orthologous proteins (BcMOY was excluded because incomplete) from *Ceratitis capitata* (MOY), *Bactrocera oleae* (BoMOY), *B. zonata* (BzMOY), *B. latifrons* (BlMOY), *B. jarvisi* (BjMOY), *B. dorsalis* (BdMOY), *B. tryoni* (BtMOY), and *Zeugodacus cucurbitae* (ZcMOY), respectively. In black bold are indicated amino acids conserved in all 8 species, in light blue those conserved between *Ceratitis* and some other species and in green those conserved in Tephritidae species other than *Ceratitis*. In red, a MOY conserved short region, named as MOY-R, in blue MOY conserved *Bactrocera* Regions, named as BMOY-R1, -R2 and-R3.

To further support the requirement of *MoY* for male sex determination, loss-of-function alleles were generated by embryo microinjections of Cas9 ribonucleoproteins^16^
targeting a predicted 70-aa MOY coding sequence (Extended Data Table 2; Extended Data Fig. 4a). Indels near the sgRNA-target site of *MoY* were detected in genomes of G0 XY larvae and G0 XY intersexes (Extended Data Fig. 5a,b). 50% of the XY individuals were transformed into phenotypic females (2/14) or intersexes (5/14) (Extended Data Fig. 4b-c, Extended Data Fig. 5b). As above, G0 XY reverted females were crossed with XX males and one of these produced female-only G1 offspring as expected, including 3 XY and 18 XX females (Supplementary Table 3; Extended Data Fig. 4d-e). Two XY G1 females had frameshift-inducing deletions (10 bp and 4 bp long, respectively) in the *MoY* open reading frame (Extended Data Fig. 5a). Similarly to RNAi-induced XY females, we observed reduced fertility and bias in favour of XX offspring, possibly due to hidden sexual mosaicism of cellular clones, reduced viability of Y-carrying eggs or problems of X-to-Y pairing during female meiosis. We concluded that *MoY* is likely a protein coding gene, that is necessary for male sex determination that acts to inhibit *Cctra* female-specific splicing and/or the establishment of its autoregulation. We also concluded that the medfly Y chromosome, either with a wild type *MoY* or carrying null *MoY* mutations, can be transmitted through XY female meiosis (Extended Data Fig. 3b).

Next, we investigated whether *MoY* is sufficient for male sex determination. A 5 kb genomic fragment, encompassing the *MoY* locus and flanking regulatory regions (Fig. 1c), was injected as a linear PCR product or as circular plasmid into either XX-only or XX/XY embryos (Fig. 2d; Extended Data Table 2). As a result, male-specific *Cctra* splicing was artificially induced in XX individuals at embryonic, larval and adult stages (Fig. 2d; Extended Data Fig. 6a-c). In injection set number 4, 75% of emerging XX flies (9/12) were either fully masculinized into fertile XX males (3/12) or partially masculinized at the phenotypic or molecular levels (6/12) (Supplementary Fig. 2, Extended Data Fig. 6b). To confirm that *MoY* is a protein coding gene we performed embryo microinjections of a recombinant his-tagged MOY protein. We recovered partial phenotypic masculinization in 19% of the emerging XX adults (6/31, Extended Data Table 2, Supplementary Fig. 3). We concluded that MOY protein controls male sex determination. In contrast to M factors that have been recently discovered in other dipterans^8-11^, *MoY* is sufficient to fully masculize XX individuals, making it a particularly attractive tool for translational research.

*Ceratitis* MOY showed no significant BLASTp hits to the NCBI non-redundant protein database, suggesting either novelty or a high sequence divergence. In contrast, tBLASTn searches of available assembled genomes, WGS or RNA-seq data of 14 Tephritidae species spanning 111 Mya^17^ identified putative MOY orthologues in 8 of them (Fig. 3a, b, Supplementary Table 8). MOY exhibits 43-60% amino acid sequence similarity to the 7 MOY orthologues (Supplementary Notes 5-12). The most conserved sequence region among the 8 MOY orthologs is located in the N-terminal portion, where a well preserved hexapeptide KXNSRT occurs (Fig. 3b, MOY-R in red). We confirmed Y-linkage of *MoY* orthologues in 4 species, namely *B. oleae, B. dorsalis, B. tryoni* and *B. jarvisi* by PCRs of male and female genomic DNA (Fig. 4a). Knock-down of *MoY* orthologs using embryonic RNAi microinjections of *B. oleae* and *B. dorsalis* embryos led to partial or full feminization of XY individuals, confirming the functional conservation of *MoY* as the male-determining Y-linked gene in other Tephritidae species (Fig. 4b; Extended Data Table 2; Extended Data Fig. 7).

**Figure 4.**
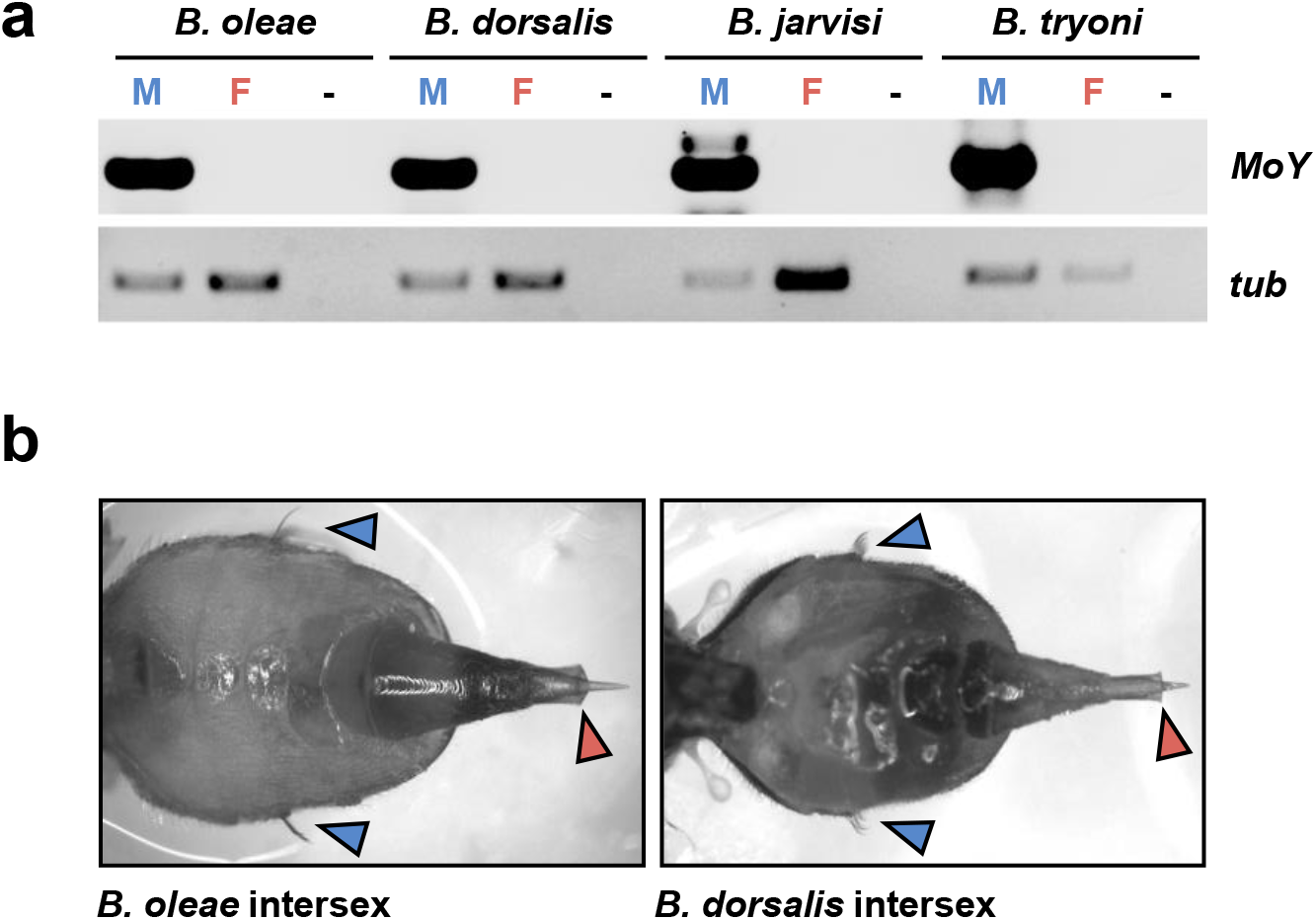
*MoY* gene is Y-linked in *Bactrocera* species species. **a**, PCR on male and female genomic DNA showing Y-linkage of *MoY* orthologues in *Bactrocera oleae* (*B. oleae*), *B. dorsalis, B. jarvisi*, and *B. tryoni*. Positive control: beta tubulin. **b**, XY intersexes of *B. oleae* and *B. dorsalis*, following embryonic *BoMoY* and *BdMoY* RNAi, which show the presence of both female-specific (ovipositor; red arrow) and male-specific characteristics (abdominal lateral bristles; blue arrows).

## Discussion

Over the course of evolution of the insect sex determining pathway, duplication and neo-functionalization of splicing-related genes generated different types of regulators which control the splicing of *transformer*^2,3,11,18,19^. In contrast, our study suggests that in Tephritidae *MoY,* being unrelated to known splicing regulators or other proteins, could have emerged *de novo*^20^, to prevent female splicing of *transformer* and the resulting establishment of its positive feedback loop. Likely, it is the intrinsic instability of a splicing-based autoregulatory loop that makes it susceptible to interference by different mechanisms and molecules^5,19,21^.

In contrast to previous studies which showed that different M factors evolved in related species (*Anopheles* genus)^8-10^ or even in the same species (*Musca domestica*)^12^, *MoY’*s structure and function appear to be highly conserved in Tephritidae species, which span over 111 Mya of divergence. To develop effective control strategies for agricultural pests and disease vectors, it will be important to understand the mechanisms by which MOY^22^ and other short novel M proteins, such as YOB^9^ and GUY1^10^ in mosquitoes, orchestrate splicing of their downstream targets in the sex determining pathway.

Despite intense control campaigns, Tephritid fruit flies are diffusing in agricultural areas world wide. For example, the olive fly from the mediterranean area has recently colonized California^23^ and the oriental fruit fly *B. dorsalis* was recently detected for the first time in Europe^24^. An emerging perspective from the presented *MoY* data is a remarkable plasticity of medfly development and possibly of other Tephritidae to perturbations of sex determination and differentiation. From a translational perspective, this plasticity makes *MoY* an excellent candidate for the future development of more efficient medfly genetic control strategies, such as *MoY*-based male-converting gene drives^25-30^. The atypical conservation of *MoY* across the Tephritid genus spanning approximately 111 Mya that we documented, makes the applied potential of *MoY* even more appealing, because technologies based on *MoY* could be easily transferable between species in a large number of agricultural pests.

**Supplementary Information** is available in the online version of the paper.

## Acknowledgements

This paper is dedicated to the memory of JN who passed away during the first phase of the project. We thank Carlos Caceres and Sohel Ahmad (FAO-IAEA, Seibersdorf, Austria), for providing *B. oleae* strain, material and methods to rear them. We thanks Bernardo Carvalho (University of Rio de Janeiro) for suggestions on bioinformatics, Jennifer Morrow (Western Sydney University) for providing *B. jarvisi* and *B. tryoni* fly samples, and Daniel Bopp (University of Zurich) for discussions, suggestions, and for a generous critical reading and editing of the manuscript We thank Serena Aceto (University of Naples Federico II) for suggestions on the phylogenetic analysis and protein alignments.

RNA and DNA Sequencing costs were covered by MDR (Swiss National Science Foundation (project numbers 31003A_143883, 310030_175841 and support from the University of Zurich, Research Priority Program Evolution in Action for SS). This research was supported by “Ricerca Dipartimentale” grants to GS and MS (Department of Biology, Federico II University of Naples, Italy, 2015-2018), by the grant STAR2013_25 to MS from University of Naples Federico II and Compagnia di San Paolo, Naples, ITALY, in the frame of Programme STAR2013 (Sostegno Territoriale alle Attività di Ricerca) and by PhD fellowships to AM and PP from PhD program in Biology (Department of Biology, University of Naples Federico II, Italy). NW was funded by the BBSRC under the research grant BB/P000843/1. ZT was supported by NIH grants AI121284 and AI123338. PAP was supported by a Rita Levi Montalcini award from the Ministry Education, University and Research (MIUR—D.M. no. 79 04.02.2014). JR was supported by Compute Canada Resource Allocation Project (WST-164-AB), and Genome Canada Genome Innovation Node (244819) grants. We would like to thank Haig Djambadzian and Anthony Bayega (McGill) for providing data and manuscript edits. This study was benefited from discussion during the research coordination meetings of several Coordination Research Projects (CRP) supported by the International Atomic Energy and particularly of the CRP on “Comparing Rearing Efficiency and Competitiveness of Sterile Male Strains Produced by Genetic, Transgenic or Symbiont-based Technologies”.

## Author Contributions

GS and JN initiated the project. GS conceived the production of XX-only embryos for mRNA-seq, high-throughput functional tests by RT-PCR (after injection of dsRNA or DNA) in early embryos to identify the M factor, by embryos injections of recombinant MOY protein, the use of FAM18 strain carrying a shorter Y, and performed the identification of *MoY* Tephritidae orthologues. MS improved the crossing scheme to produce XX-only embryos. GS, MDR, MS, ZT, BH, AK, JN, PAP, NW designed and performed bioinformatics analyses, with contributions from AM and PP. ZT, BH performed preliminary CQ analysis. MS, VP and AM prepared RNA and DNA for sequencing. MDR performed mRNA sequencing, DE and CQ analyses, PacBio FAM18 sequencing, and Canu assembly together with SS. MS performed the medfly 4-8h embryonic transcriptome assembly and the Tephritidae species transcriptome assemblies. MS developed the local web-tool with graphical interface utilized for all the BLAST searches of this paper. AM and GS selected *MoY* as a first M candidate from the list of putative male-specific transcripts by MDR and the list of novel putative male-specific transcripts by ZT and BH (Supplementary Note 2). AM performed eRNAi, CRISPR/Cas9, and *MoY* DNA injections and RT-PCR analyses demonstrating *MoY* function. PP, AG, MAG, FF, DI and MMP maintained the strains, performed crosses and DNA/RNA molecular analyses. LV and AR purified recombinant MOY protein and performed structural/similarity analyses to protein databases. PP performed MOY protein embryos injections. FM and MD performed *in situ* hybridization of *MoY*. KM, JR, KTT, MEG performed qRT-PCR analysis of *MoY* in *C. capitata* and *MoY* expression analysis in *Bactrocera oleae*. KM and JR provided *Bactrocera oleae* genome assembly data. AM, FS, PK and KB performed *MoY* RNAi on *MoY* orthologues of *Bactrocera oleae* and *Bactrocera dorsalis.* PK and KB performed molecular analysis on reverted XY females in these two species. GS, AM, MS and PAP prepared the figures. All of the authors discussed the data. PAP, SMM, EG and LV provided essential reagents. GS wrote the manuscript with intellectual input from all authors, especially PAP, MDR, MS, and LV. AM and MS contributed equally to the work. GS supervised the project.

## Author Information

The 12 Kb *MoY* sequence region has been deposited in GenBank under accession number MK330842. Reprints and permissions information is available at. The authors declare no competing financial interests. Readers are welcome to comment on the online version of the paper. Correspondence and requests for materials should be addressed to GS, MDR, or PAP.

## Methods

### Insects rearing and strains

The following medfly strains were used in this study: 1) the wild-type strain *Benakeion* developed by P. A. Mourikis (Benakeion Institute of Phytopathology, Athens, Greece); 2) the Fam18 strain^6^ (see below); 3) the *Vienna8* strain^12^, carrying a Y chromosome marked with translocated wild type allele of a *white pupae* mutation. The *C. capitata Benakeion* strain was reared in standard laboratory conditions at 25 °C, 70% relative humidity and 12:12 h light–dark regimen. Adult flies were fed yeast/sucrose powder (1:2). Eggs were collected in trays filled with distilled water and transferred to larval food (Piccioni lab, Italy) after hatching. Pupae were collected and stored in dishes until eclosion.

The *Bactrocera dorsalis* Saraburi strain originated from Thailand and was maintained at the FAO/IAEA Insect Pest Control Laboratory (IPCL, Seibersdorf, Austria) for 84 generations before use in these experiments. Flies were reared at 25 ± 1 °C, 60 ± 5% RH and a photoperiod of 12h light : 12h dark, and fed with a standard laboratory adult diet containing sugar and hydrolyzed yeast in a 3:1 ratio and water *ad libitum*. Eggs were collected and transferred to larval medium containing 28% wheat bran, 7% brewer’s yeast, 13% sugar, 0.21% sodium benzoate, 0.21% nipagin, 1% HCl, 50% water.

The *Bactrocera oleae* Greece-Lab strain originated from a stock in the Department of Biology, ‘Demokritos’ Nuclear Research Centre, Athens, Greece, and was maintained at the FAO/IAEA Insect Pest Control Laboratory IPCL for 177 generations. Flies were reared at 25 ± 1 °C, 60 ± 5% RH, a photoperiod of 14h light : 10h dark, and fed with a standard laboratory adult diet consisting of 75% sugar, 19% hydrolyzed yeast and 6% egg yolk powder. Eggs were collected and transferred to larval diet containing 550 mL water, 20 mL Extra Virgin Olive Oil, 7,5 mL Tween® 80 emulsifier, 0.5 gr potassium sorbate, 2 gr nipagin, 20 gr sugar, 75 gr brewer’s yeast, 30 gr soy hydrolysate, 4.5 mL HCl 36% and 275 gr cellulose powder.

### Production of XX-only embryos

Embryonic RNAi targeting *Cctra* was used to produce male-only progeny including XX fertile reverted males, as previously described^5^. 40 males were individually crossed 3 wild type virgin females each. 4-8h AEL embryos were collected in 40 separated samples for RNA extractions and RT-PCR assays using Y-specific marker (*Cclap*). 22 RNA samples corresponded to XX-only progeny from individual crosses with XX fertile males and were pooled in 3 sets for sequencing.

### RNA sequencing, *de novo* transcriptome assembly and differential gene expression

Total RNA was prepared from 4-8 h AEL (when male sex determination occurs^31^) *C. capitata Benakeion* strain embryos of mixed sexes (XX/XY) and of female-only sex (XX; see above), in three biological replicates each, with TRIzol reagent (Thermo Fisher Scientific) according to the manufacturer’s protocol. The integrity and purity of extracted total RNA were assessed using the NanoDrop 2000c (Thermo Fisher) and the Agilent 2100 Bioanalyzer with RNA 6000 Nano kit (Agilent, Santa Clara, CA, US). All RNA samples had a A260/280 ratio higher than 2.1 and a RIN value higher than 7.8. Total RNAs were depleted of ribosomal RNA with Ribo-Zero rRNA Removal kit (Epicentre). Libraries for sequencing, with an average insert size of 300 bp, were generated with the NEBNext Ultra Directional RNA Library Prep Kit for Illumina (New England Biolabs). Illumina stranded and paired-end (PE) sequencing was performed at NXT-GNT (Belgium) yielding a total of 85,832,487 XY/XX and 94,085,461 XX-only PE 100 bp-long reads (BioProject PRJNA434819 in SRA archive). A transcript catalogue was *de novo* produced using the whole 4-8 hrs Illumina read data set, concatenated into two paired FASTQ files and the Trinity assembler^32,33^, run with default parameters and -SS_lib_type RF -jaccard_clip -normalize_reads flags set, after adapter removal and quality trimming using Trimmomatic v.0.33^34^ and mitochondrial and ribosomal reads depletion, as described^35^. Assembly statistics are presented in Supplementary Table 4. Transcript-level quantifications for each sample were done against this catalogue using Kallisto^36^ (Supplementary Table 5). Differential gene expression was performed using edgeR^37^, with cut-off values of FDR < 0.05 and logFC > 0 (Supplementary Table 6).

### *Fam18* strain description

Due to the lower complexity of its shorter Y chromosome, the *Fam18* strain (kindly provided by Dr Franz, FAO/IAEA, Austria), was used for PacBio sequencing of male genomic DNA. This strain originated from a study by Willoefth and Franz^6^, who used cytogenetic approaches, including *in situ* hybridization with Y-specific repetitive probes^38^ and Y-autosome translocations, to map the male-determining factor region on the long arm of the *C. capitata* Y chromosome, toward the centromere. This region represents approximately 15% of the entire medfly Y chromosome. In *Fam18*, males carry an internal deletion of the long arm of the Y chromosome but not Y-autosomal translocations, and this Y chromosome lacks a segment between the M region and a point toward the tip (Franz, G., FAO-IAEA Pest Control Unit, person. comm. to GS, 2010). To verify that in the *Fam18* males there is a reduced complexity of the Y chromosome due to the large deletion of its long arm, we mapped, using the Bowtie aligner^39^, the *Fam18* males and females Illumina reads against four Y-specific repetitive elements, known to be widely distributed all over the medfly Y chromosome^6,38^. *Fam18* read counts are significantly lower for three out of four Y-linked repetitive elements compared to the medfly ISPRA standard strain read counts (Supplementary Table 7). Conversely, the read counts for the control gene are comparable between the two strains.

### Illumina short read WGS of *C. capitata Fam18* strain

Five adult males and five adult females from the *C. capitata Fam18* strain were collected and genomic DNA was extracted from the two sexed pools separately, as described below^40^. Each pool was sequenced with 1 lane of Illumina HiSeq2500 resulting in 243,152,822 male and 248,886,022 female 2×126-bp PE reads, respectively (BioProject PRJNA435549 in SRA archive).

### Illumina short read WGS, trimming, error-correction and assembly

The Illumina raw short-read data for the males had approximately 126X coverage (assuming a genome size of 485Mbp). Trimmomatic v0.35^34^ removed the Illumina adapters in paired-end mode, trimmed by quality values in maxInfo mode with target_length=0 and strictness=0.4 parameters, and selected for a remaining read length of at least 50 bases. The resulting trimmed reads were used for all purposes except for the error correction of the PacBio reads. The trimmed reads had a coverage of 62X for the first and 59X for the second read.

For the PacBio correction, we first corrected the trimmed short reads with BFC^41^ (r181) using k-mer length=33 and genome size=485Mb as parameters. Meraculous^42^ assembled those corrected short reads into unitigs with genome size=485Mb, num_prefix_blocks=8, and automatic k-mer length identification. To determine the library parameters for the corrected short reads, we mapped the reads to the Ccap01172013 genome from Baylor College^7^ using bowtie2^43^ v2.2.5 with a maximum insert length of 1000. This resulted in a mean insert size of 364 bp with a Gaussian standard deviation of 79 bp. The average read length of the corrected reads was 121 bp. The corrected reads and the unitigs were then used to correct the PacBio reads.

### PacBio Whole Genome Sequencing (WGS) of *C. capitata Fam18* strain

Five adult males from the *C. capitata Fam18* strain were collected and genomic DNA was extracted using the Holmes-Bonner protocol^40^. Extracted genomic DNA was resuspended in 10 mM Tris-HCl, pH 8.5, and its integrity and purity were assessed using the NanoDrop 2000c (Thermo Fisher Scientific) and 1% agarose gel electrophoresis, resulting in A260/280 ratio of 1.8. Genomic DNA was sheared by g-TUBE (Covaris) and a BluePippin instrument (Sage Science) was used to select fragments from 15-20 Kb in length. A large insert library was prepared using Pacific Biosciences recommended protocols and P6-C4 reagents. The resulting sample was sequenced at the Functional Genomics Centre Zurich (Switzerland) with 20 SMRT cells on a Pacific Biosciences RSII instrument. The PacBio SMRT cell reads have been submitted to the NCBI SRA archive with the following accession number: BioProject PRJNA435534.

### PacBio WGS error-correction and assembly

PacBio DNA sequencing using P6-C4 chemistry for 20 SMRT cells was conducted; extraction of sequences into FASTQ files was done by Dextractor (https://github.com/thegenemyers/DEXTRACTOR) with a minimum sequence quality parameter set to 400 produced ∼32X genome coverage (assuming a genome size of 485Mbp) with an average read length of 5,526 bp and a N50 length of 8,656 bp, and a maximum length of 68,782 bp. We combined all SMRT cells and then split the data into chunks of 400 Mb for cluster processing. The error correction was done by Proovread^44^ using the corrected first reads from the PE short read data and the unitigs. We did not use the second reads as they were of lower quality and the additional coverage would have slowed down the error correction without improving the result. The only additional parameter set was coverage=62. This resulted in a total of ∼19X corrected sequences with an average read length of 2,855 bp, N50 length of 4,695 bp, and maximum length of 33,053 bp. This set of corrected reads was used to build a *de novo Fam18* genome assembly using Canu v1.0 with -pacbio-corrected set and genomeSize=0.45g. This resulted in 12,165 assembled contigs with a total length of 496 Mb, an average read length of 40,769 bp, N50 length of 62,964 bp, and maximum length of 1.0 Mbp.

### Bioinformatics for *MoY* gene structure

BLASTn search of the PacBio Canu assembly, using medfly *C. capitata MoY* 681 nt-long transcript sequence (including 5’ and 3’ UTRs) as DNA query (Genbank acc. num. MK165756), identified a highly similar corresponding genomic sequence (95% identity). The *MoY* putative coding region of the transcript is 99% identical in the PacBio genomic sequence, with only 2 SNPs, with the second inducing a conservative amino acid substitution at position 63 (I->M) (Genbank acc. num. MK165755).

### Probe preparation and labelling

1.5 kb fragment of *MoY* plasmid was amplified by PCR using the primers *New_MoY_F1*/*New_MoY_R*. The reaction with a final volume of 20 μl contained 22 ng of plasmid DNA, 0.2 mM of each dNTP, 1 μM of each primer, 1× Ex *Taq* buffer, and 1U Ex *Taq* DNA Polymerase (TaKaRa, Otsu, Japan). The PCR product was purified using Wizard SV Gel and PCR Clean-Up System (Promega, Madison, WI, USA) according to manufacturer instructions.

Probe labelling was performed using an improved nick translation procedure^45^ with some modifications. The modified 40 µL reaction contained 500 ng of unlabeled DNA, 50 µM of each dATP, dCTP, and dGTP, 10 µM of dTTP, 20 µM digoxigenin-11-UTP (Roche Diagnostics, Mannheim, Germany), 1× nick translation buffer (50 mM Tris-HCl pH 7.5, 5 mM MgCl2, 0.005% BSA), 10 mM ß-mercaptoethanol, 1.25×10-^4^ U/µl DNase I and 1 U/µl DNA polymerase I (both ThermoFisher Scientific, Waltham, MA, USA). The reaction was incubated at 15 °C for 40 min and then purified using an illustra Sephadex G-50 column (GE Healthcare, Buckinghamshire, UK).

### Chromosome preparation

Spread preparations of mitotic chromosomes were made from the brain (cerebral ganglia) of third instar larvae following a previously described method^46^ with slight modifications. Briefly, the brain was dissected in a physiological solution, transferred into hypotonic solution (0.075 M KCl) for 10 min, and then fixed for 15 min in freshly prepared Carnoy’s fixative (ethanol:chloroform:acetic acid, 6:3:1). The fixed tissue was spread in a drop of 60% acetic acid on the slide at 45 °C using a heating plate. Then the preparations were passed through a graded ethanol series (70%, 80%, and 100%, 30 s each), air dried, and stored at −20 °C. Before further use, the preparations were again dehydrated in the ethanol series, immediately after removal from the freezer.

### Fluorescence *in situ* hybridization with tyramide signal amplification (TSA-FISH)

The TSA-FISH was performed according to published protocols^46,47^ with some modifications. Slides were pre-treated with 200 µg/mL RNase A in 2× SSC at 37 °C for 1 h, with 0.01 M HCl at 37 °C for 20 min, 1% H_2_O_2_ in 2× SSC at room temperature (RT) for 30 min, and 5× Denhardt’s solution (0.1% Ficoll 400, 0.1% polyvinylpyrrolidone, and 0.1% bovine serum albumin in ultrapure water) at 37 °C for 30 min. The probe cocktail containing 50% deionized formamide, 10% dextran sulphate, and 0.5 ng/µL of the probe in 2× SSC was applied to chromosome preparation, covered with cover slip and denatured at 70 °C for 8 min. Hybridization was carried out at 37 °C overnight.

After hybridization, the slides were washed three times in 50% formamide in 2× SSC at 46 °C for 5 min, three times in 2× SSC at 46 °C for 5 min, three times in 0.1× SSC at 62 °C for 5 min, and in TNT buffer (0.1 M Tris-HCl pH 7.5, 0.15 M NaCl, 0.05% Tween-20) at RT for 5 min. The slides were blocked in TNB buffer (0.1 M Tris-HCl pH 7.5, 0.15 M NaCl, 0.5% Blocking Reagent; PerkinElmer, Waltham, MA, USA) at RT for 45 min. The probe was detected by anti-digoxigenin-POD (Roche Diagnostics) diluted 1:20 in TNB buffer. Tyramide amplification was carried out using the TSA™ Plus Fluorescein System (PerkinElmer) according to the manufacturer’s instructions. The slides were incubated with tyramide working solution at RT for 20 min and then washed as described previously^46^. Chromosomes were counterstained with 0.5 µg/mL DAPI (4′,6-diamidino-2-phenylindole; Sigma-Aldrich, St. Louis, MO, USA) in 1× PBS at RT for 15 min. Finally, the slides were mounted in antifade based on DABCO (1,4-diazabicyclo(2.2.2)-octane; Sigma-Aldrich). Chromosome preparations were observed in a Zeiss Axioplan 2 microscope (Carl Zeiss, Jena, Germany). Black-and-white images were captured separately for each fluorescent dye with a monochrome CCD camera XM10 using cellSens Standard software version 1.9 (both Olympus Europa Holding, Hamburg, Germany). The images were pseudo-coloured and merged using Adobe Photoshop CS5 (Adobe Systems, San Jose, CA, USA).

### Evaluation of *MoY* expression via qRT-PCR: RNA isolation, cDNA synthesis and primer design

For the validation of the *MoY* gene expression, RNA was extracted from several developmental stages and tissues of the medfly. A pool of ∼50 eggs were removed from the incubator at different time intervals throughout embryonic development. Each embryonic sample refers to the time after oviposition, at which eggs were collected in Extrazol (Nanogen Advanced Diagnostics, Turin, Italy). In total, 10 different embryonic time points, i.e. 1h, 2h, 3h, 4h, 5h, 6h, 7h, 8h, 15h and 48h, were examined in three biological replicates. Regarding the other developmental stages, a pool of 10 per sample was collected for 1^st^ and 3^rd^ instar larvae respectively, whereas a pool of 5 per sample was collected for pupae, female adults, male adults, as well as for testes. These aforementioned samples were examined in either three or two biological replicates (in the case of male adult tissues).

Following RNA extraction, genomic DNA was removed using a DNase treatment with 1.0 unit of TURBO™ DNase (Invitrogen, CA) according to manufacturer’s instructions. The total amount of DNA-free RNA obtained from each tissue was reverse transcribed using the PrimeScript RT reagent Kit (TaKaRa, Japan) according to manufacturer’s instructions. Reverse transcription was conducted at 37°C for 15 min and 85°C for 5 min. The resulting cDNA was used in the subsequent qPCR reactions. Specific primers were designed by IDT primer-designing software PrimerQuest on the Integrated DNA Technologies (IDT) website (http://www.idtdna.com) for the amplification of *CcMoY* (**Supplementary Methods** Table 1). A quantitative Real-time PCR (qRT-PCR) approach was performed to analyze changes in *MoY* transcription. The expression levels were obtained relatively to the suitable housekeeping gene based on the evaluating reference genes list of each species^48^. The *β-tub* gene was used to normalize expression in eggs, whereas *Rpl19* for larvae, pupae, male and female adults respectively (Supplementary Methods Table 2). The qRT-PCR cycling conditions were: polymerase activation and DNA denaturation step at 95 °C for 3 min, followed by 50 cycles of 10 s at 95°C, 20 s at the annealing temperature, and 30 s at 72°C. Each qPCR run was followed by a melting curve analysis (gradual increase of temperature between 55 and 95°C) to examine the specificity of each reaction. For all primers, a single gene specific peak was observed, confirming the absence of non-specific products or primer dimers. Each reaction contained 5μl from a dilution 1:10 of the cDNA template, 1X KAPA SYBR FAST Universal qPCR Master Mix (Kapa Biosystems) and 300nM of each primer, to a final volume of 15μl. The reactions were carried out on Bio-Rad Real-Time thermal cycler CFX96 (Bio-Rad, Hercules, CA, USA) and data analysis was performed using the CFX Manager™ software.

### Embryonic RNA interference

*CcMoY, BdMoY, and BoMoY* fragments were PCR amplified using genomic DNA from a male adult of *C. capitata, B. dorsalis* and *B. oleae* (List of primers in Supplementary Methods Table 1), introducing a T7 promoter sequence at each end. PCR was performed with Phusion^®^ High-Fidelity DNA Polymerase (New England Biolabs) according to manufacturer instructions. PCR product was purified with a phenol/chloroform purification and eluted in ddH2O for a final concentration of 1 μg/μL. *In vitro* transcription of *MoY* dsRNA for all the species was obtained using Ambion MEGAscript^®^ RNAi kit T7 RNA polymerase, following manufacturer instructions. Final product was checked on 1% agarose gel electrophoresis and quantified on NanoDrop 2000c (Thermo Fisher Scientific). The dsRNA was eluted from the purification column using elution buffer, containing 5 mM KCl and 0.1 mM PO4, pre-heated at 98 °C, to get a final dsRNA yield of 1.5 μg/μL.

Embryos were collected 45 min AEL, placed on double-stick tape, hand-dechorionated, dehydrated in a Petri dish containing calcium chloride, covered with Halocarbon oil 27 (Sigma, H8773) for the three species. A solution of 1 μg/μL dsRNA was loaded inside the needle using a P10 with Eppendorf Microloader^®^ capillary tips. For the microinjections in *Ceratitis capitata*, an inverted contrast microscope Leica DM IRB^®^ was used. The needles were obtained from Boron-Silicate cylinders with a diameter of 0.5 mm pulled with a puller machine (Narishige). The injections were conducted in the posterior pole of the embryos, with the needle entering the embryo for about 30-50% of its length, and the pressure was generated using an oil pump. Injections in *Bactrocera spp.* were done into the posterior end of the eggs using XenoWorks® Digital Microinjector and Micromanipulator System (Sutter Instrument) and Quartz needles (#QF100-70-10), pulled in P-2000 Laser-Based Micropipette Puller (Sutter Instrument).

### RNA extractions and RT-PCR on *C. capitata*

Total RNA was extracted from pools of embryos, larvae, and intersexes, and male and female adults using TRIzol Reagent (Thermo Fisher Scientific) following manufacturer instructions. Oligo-dT-primed cDNA was prepared from DNase I-treated total RNA using EuroScript^®^ m-MLV reverse transcriptase (Euroclone^®^). RT-PCR expression analysis was performed using primers described in Supplementary Methods Table 1.

### Splicing of *Bdtra* and *Botra*

Total mRNA was extracted from eggs 12 hours after injection and 3rd instar injected larvae using TRIzol Reagent (Thermo Fisher Scientific) and treated with TURBO™ DNase (Invitrogen) according to manufacturer’s instructions. cDNA was synthesized using SuperScript™ III First-Strand Synthesis System (18080051) following manufacturer’s instructions and PCR amplification of *tra* was performed using Qiagen Taq PCR Master Mix Kit (201443) according to manufacturer’s instructions. RT-PCR expression analysis was performed using the primers reported in Supplementary Methods Table 1.

### Molecular karyotyping of *Bactrocera* adults

DNA was extracted using EXTRACTME DNA TISSUE kit (EM03) following manufacturer’s instructions. PCR was performed using Qiagen Taq PCR Master Mix Kit (201443) according to manufacturer instructions. The primers for the amplification of the Y-specific fragment were *BdMoY-F* and *BdMoY-R* for *B. dorsalis; BoMoY-F* and *BoMoY-R* for *B. oleae.* PCR analysis was performed using primers described in Supplementary Methods Table 1.

### CRISPR and RNP complex assembly in *C. capitata*

sgRNA was designed using CHOPCHOP^49^. Following, production of template for sgRNA was performed, with minor modifications, as described^16^ with CRISPR-*MoY-F* and the invariant reverse primer (PAGE-purified, Life Technologies) (Supplementary Methods Table 1). sgRNA was synthesized according to instructions of the Megascript^®^ T7 kit (Ambion) with 1 μg of template and a 5’ flanking T7 promoter as starting material. After RNA synthesis, template was removed by incubating with TurboDNase^®^ (Ambion) for 15 min at 37 °C. Cas9 was expressed as a His-tagged protein and purified from bacteria as described^16^. Prior to the injection, the RNP complex was prepared by mixing 1.8µg of purified Cas9 protein with approximately 200 ng of sgRNA in a 5?μL volume containing 300?mM KCl^16^. The mix was incubated for 10 min at 37 °C. A glass needle was filled with the pre-loaded sgRNA-Cas9 mix and the injection was performed into the posterior end of embryos collected 45 min AEL as described for RNA interference in *C. capitata*. 48 hours after the injections, the slides are inspected for hatched larvae under a stereomicroscope. Hatched larvae are transferred to Petri dishes containing larval food.

### Amplification and cloning of the 5 Kb fragment of *CcMoY*

PCR on the 5 Kb fragment of *CcMoY* was carried out with the LongAmp® Taq DNA Polymerase following manufacturer’s instructions in a final volume of 100 μL using the primers *Gen_CcMoY_F* and *Gen_CcMoY_R* (Supplementary Methods Table 1). The 5 kb *CcMoY* fragment was then cloned in the pGEM®-T Easy vector (Promega) following manufacturer’s instructions. The PCR product and the plasmid with the 5 Kb insert were purified with phenol:chloroform (1:1), precipitated in ethanol and eluted with injection buffer (containing 5 mM KCl and 0.1 mM PO4) for a final concentration of 1 μg/μL to be injected in *C.capitata* embryos.

### MOY protein expression, purification and characterization

The *MoY* gene was amplified using a male adult *C. capitata* genomic DNA as template with two synthetic primers carrying the *NcoI* and *XhoI* restriction sites: *CcOrf2fw (NcoI)/ CcOrf2rv (XhoI)*. The fragment was then cloned into the pETM-13 expression vector. The resulting gene contained an extra C-terminal tag of six histidine residues. The MOY protein was then heterologously expressed in *Escherichia coli* BL21 (DE3) cells. MOY was purified from inclusion bodies after removing the soluble proteins by sonicating the bacterial resuspension in a native buffer. The pellet was then resuspended in a denaturing buffer containing 8 M Urea, 50 mM Tris-HCl and 200 mM NaCl (pH 8.0) and left at room temperature for 2 hours to facilitate its solubilization. After the centrifugation, the supernatant was loaded on a Ni-NTA resin (Qiagen) equilibrated with the denaturing buffer. The protein was washed with ten volumes of this buffer and then was eluted with a solution containing a high concentration of imidazole (150-300 mM pH 8.0). The homogeneity of the eluted protein was evaluated by SDS–PAGE analysis. MOY protein was refolded in solution containing 20 mM ethanolamine and 0.5 M L-arginine (pH 9.0-10.0) using rapid dilution (1:50 v/v) with three additions of the same volume over a 24h period. The protein was concentrated using Amicon centrifugal devices (Millipore) up to a concentration of 70 µg/mL. The molar mass of the refolded MOY was determined by mass spectrometer (ESI-TOF) (9250.28 Da) in agreement with that computed from the tagged sequence (9251.7 Da).

### *Bactrocera* spp. transcriptome assemblies and *MoY* orthologs search

Illumina RNA-seq data for 14 *Bactrocera* species were downloaded from NCBI SRA archive and *de novo* assemblies were produced using Trinity assembler^32^ with default parameters and-normalize_reads and -jaccard_clip flags set. Assembly statistics and accession numbers of the utilized data set are listed in Supplementary Table 8.

tBLASTn search of the *B. oleae* transcriptome, using MOY as protein query, identified a transcript encoding a putative protein with 36% identity over a 64 aa-long region (*BoMoY*, 70 aa; MK165746) (Supplementary Notes 5-12). TBLASTN search, using BoMOY on *B. oleae* NCBI WGS database, failed to find the full-length corresponding *BoMoY* gene, but revealed the presence of 4 genomic sequences (Sequence IDs: JXPT01043932.1; LGAM01008500.1; LGAM01009849.1; LGAM01000393.1) encoding for truncated 15-41 aa-long *BoMoY* paralogous sequences and showing 37%-100% aa identity to BoMOY protein. The identified *B. oleae MoY* ortholog was utilized to search by tBLASTn for *MoY* orthologues in the other thirteen assembled *Bactrocera* transcriptomes. *MoY* orthologs were found only in *B. jarvisi* (MK165748) and *B. zonata* (MK165752). As we reasoned that *MoY* transcripts could be underrepresented in assembled transcriptomes for both technical limitations and low gene expression levels, we searched for *MoY* orthologs by tBLASTn searches at NCBI on SRA and WGS databases. Sequences encoding full-length *MoY* orthologs were found in both *B. tryoni* (JHQJ01009763.1) and *B. latifrons*, showing 63% aa identity with *BoMoY* (MIMC01001452.1). tBLASTn analyses revealed also the presence in *B. tryoni* of a shorter *MoY* paralogue (50 aa long) in the same genomic contig, 4 Kb apart from the first *BtMoY* copy (70 aa long). The two *BtMoY* paralogues showed 90% protein identity over a 50 aa-long region. Similarly, in *B. latifrons* we have found a truncated paralogue *BlMoY* gene in a different genomic contig (MIMC01001198.1), encoding for truncated 36 aa-long *BlMoY* sequence, with 78% identity to *BlMoY* and 63% to *BoMoY*. Finally, by tBLASTn search we identified a SRA read from *B. dorsalis* transcriptome analysis data (SRR316210) encoding part of the putative *BdMoY*, corresponding to the amino-terminus. We performed PCR on cDNA from 5-8 hours old embryonic RNA using primers *Bd_MoY+*/*Bd_MoY-*designed on the corresponding paired Illumina reads (Supplementary Note 8), leading to the isolation of the *BdMoY* full-length coding region.

We validated the *MoY* male-specific Y-linked genomic localization by amplification on genomic DNA of *B. oleae, B. dorsalis, B. tryoni* and *B. jarvisi* using the specific pair of primers reported in Supplementary Methods Table 1. The following list comprises the Genbank accession numbers of the 8 putative MOY orthologous protein sequences: BoMOY-A MK165746, BoMOY-B MK165747, BjMOY MK165748, BdMOY MK165749, BtMOY MK165750, BlMOY MK165751, BzMOY MK165752, RzMOY MK165753, ZcMOY MK165754.

