## Supplementary notes for "*Maleness-on-the-Y* (*MoY*) orchestrates male sex determination in major agricultural fruit fly pests"

### Table of Content

**Supplementary Note 1 | The screening and identification of *MoY*, a single copy intron-less gene. *MoY* has some weakly related sequences in the medfly embryonic transcriptome and genome.**.. 2

**Supplementary Note 1 | The screening and identification of *MoY*, a single copy intron-less gene. *MoY* has some weakly related sequences in the medfly embryonic transcriptome and genome.**

We constructed RNA-seq libraries from embryos 4-8 h after eggs laying (AEL) from both mixed (XX/XY) embryos and female-only (XX) embryos (both from *Benakeion* strain)[12,19]. We then generated a new *de-novo* transcriptome assembly and inferred differential expression and putative chromosomal positions using male and female genomic DNA data from the *Fam18* strain. By filtering for genes predicted to be on the Y and expressed in the mixed XX/XY embryonic transcriptome but absent in the XX female transcriptome, we selected 19 candidate transcripts as XY-specific (out of 96 with XY-biased expression), corresponding to 10 distinct transcriptional units (Fig. 1a; Extended Data Table 2). Of these 19 transcripts, 7 were classified as *M* candidates of minor priority because they did not map to the *Fam18* PacBio male genome assembly and hence there were most likely linked to the Y region deleted in this strain (Extended Data Table 2). 11 out of the remaining 12 transcripts were considered of minor priority as they showed similarity to multiple long paralogous sequences in the *Fam18* genome, as most of the *M* candidates selected in the first bioinformatic screening.

However, we still investigated if any of these 13 transcribed sequences showed similarity by BLASTn to XY but not XX embryonic transcripts of the related Tephritidae species, *B. oleae*, and selected three transcripts (Extended Data Table 2). We focused on one of these 3 (DN40292_c0_g3_i1) because it is present in only one scaffold in the *Fam18* male genome, it shows sequence identity to unique transcript sequences in the XX/XY but not XX 4-8 h AEL *Ceratitis* embryonic transcriptome and, surprisingly, it corresponds to a transcript previously identified with the first analysis (Extended Data Table 1: *Corvus*). Functional analysis (see below) confirmed that this gene corresponds to the medfly male-determining factor and we thus named it *Maleness-on-the-Y* (*MoY*).

The other 2 out of these 3 transcripts (TRINITY_DN40516_c0_g2_i1 and TRINITY_DN40142_c1_g1_i5) show by BLASTn highly similar sequences present in many different Canu assembly FAM18 contigs, as shown below.

**>TRINITY_DN40516_c0_g2_i1**

```
GTTTTAAATTGAGAAATATTTTTGATTTGCCTACAAAATTGCAAACGTTTGGCATTTGGAAAGGGAGATTACAAGGGGAAATATGTATAGGCTT
TCATTTTAGGGTAGCGAGAATGAAATATCATATGACATTTTTTCTTCGTTTTGTGATTATTTTCACCACTGCCGATTATTCGAGATTATTCGAG
TCTTCTCTAAATTATCTGTTCTGCTAGTATGTGTCCCGTTATGTAGCCAGTATTTCCATAATATTCTAACTCGAGTCCTTTCTAAATTGTCTGT
```

```
TGATTGTATTTCTTGATGAAATATTTCAAATATCCTTATCCAATTTTATTTAAATAAACGCTTAAATCACAAGATAATAATACTGATCAGTAGT
GCTACAAACAGTATAAGATTTTTTAATTGGCATTGCGTTAATGTCTTGATCGTGGATGTGATAACACATCTTACGTATTAGGTTAGTTTATTGA
GACAGAAAACAGAGTAGAGCAGATTAAGTTAGAGTAGAATAGAACAGCTGTAAAGAAACTGTAGAGCAAGTTAGAGTAACTCGGAGCAGAAAGT
CTCATAAAACACGTTCTAGGTAGAAATTCTCAAAACAGTTGAATGTCCGAATTTTCAATTGACGAATATCAAATGTTTTAGATATATGGATTTC
TGTACATATACCATTTTTGGAAAATAAAAAACGGATGAATATATTATCACCGAAATCCTGGAAGAATTTCTATTGTAAATTCTTCAGACTCAGA
TAATTCGCTTTTATCCTCATCAGATGAAGAGTTTATGATGCATTTTCGTTGCTCTATTTCTGCTTCGAGAAGACTCATGAAACAAAGACTAGAG
CAAGTAAGAGCAAATAAGTGTCATTAAACATAATGTTGTGAGGTAGAGCAACCTAGCTTAACCTAGAGAAGCTCAATAAACAAACCTATTAAGT
GTCAACGAGTAGTTCACATAATTATCTCCTATAATTATCTAATTAATTTTTTTGAACAAATTCACGACGAAGAAACCTTATCGCTTACACTTCG
ATAACCGTTCGGAATTCTTTTCAAAAAACCAGCATGGATCTACAAAACTGCAAATGCTATGTGGAGGCAAGTGAATCTATTTGAGAAATGTTTT
CACCGATTTATTTAAGTGAATAACAATATAAAAATATCTCCATCGCCTTAAACAAACTTAGCGTAGCGTGAATAGCTGCATCAAGGAAGAACGC
TATGTTTGATTTTTTCTCCGTTCTGTGAGTATTTTCACTACTTTTAATTTGCCTGTAAAATTGCAGCCCTCAAGCAGTTGCAAAATGTAACTAG
AAGGGGACCTTGGTTCCCTTTATAGAGTAGCTCAAATGGCAAAAACAAGGTAGTATGATATGCTTGAAGAAGAACTTGTTTCTAATGGAGTATA
CGTTTGAGACACTTAATCGCATGCAGTTTCGCGAAATTAGTTGAAAAAATTCAGGATGAAAACCATGAAAAGCGATTTCTGAGTATTTTCGATA
ATCATTTAAAAATAGTAAAAGGAGTTTTGAGATATATCATTTCCATTTTTCTTTCATATATAAACATTTTGGGAAATATTTTTAATTCGCCTAT
AAAATTGCAGACTTGTCTCATTTATCGAAGGAAATTAGAAGCGAACAGGTATATAGGTTTTCTTTATAGAGCAGCGTGAGTGGCTGCAACAAGG
AAGTACGATATGTTTGAAGTAGACATTTTTTCTTCGTTTTTTTTGTGATTATTTGCACTAAT
```

**BLASTn in PacBio Canu assembly:**

**Query=** TRINITY_DN40516_c0_g2_i1 Length=1754

```
                                                                      Score     E
Sequences producing significant alignments:                          (Bits)  Value

  tig00011793 len=16988 reads=186 covStat=-59.19 gappedBases=no c...  3122    0.0
  tig00019373 len=35768 reads=21 covStat=127.90 gappedBases=no cl...  3112    0.0
  tig00019372 len=8510 reads=39 covStat=8.13 gappedBases=no class...  3077    0.0
  tig00021013 len=6234 reads=8 covStat=15.05 gappedBases=no class...  3068    0.0
  tig00020139 len=27776 reads=73 covStat=68.73 gappedBases=no cla...  2984    0.0
  tig00019817 len=10935 reads=41 covStat=18.09 gappedBases=no cla...  2946    0.0
  tig00019818 len=10394 reads=2 covStat=16.61 gappedBases=no clas...  2778    0.0
  tig00007607 len=12605 reads=4 covStat=37.89 gappedBases=no clas...  1651    0.0
  tig00014668 len=5823 reads=4 covStat=10.26 gappedBases=no class...  1631    0.0
  tig00013010 len=12420 reads=10 covStat=26.18 gappedBases=no cla...  1335    0.0
  tig00020818 len=11433 reads=10 covStat=25.59 gappedBases=no cla...  1330    0.0
  tig00011001 len=12237 reads=39 covStat=22.27 gappedBases=no cla...  1252    0.0
  tig00010055 len=12860 reads=18 covStat=39.45 gappedBases=no cla...  1234    0.0
  tig00020587 len=14310 reads=42 covStat=35.74 gappedBases=no cla...  1184    0.0
  tig00019374 len=7827 reads=37 covStat=6.07 gappedBases=no class...  1110    0.0
  tig00010611 len=8603 reads=2 covStat=13.53 gappedBases=no class...  1086    0.0
  tig00019375 len=16546 reads=64 covStat=23.92 gappedBases=no cla...  688     0.0
  tig00007475 len=25962 reads=38 covStat=86.12 gappedBases=no cla...  482     4e-134
  tig00000448 len=193066 reads=608 covStat=516.65 gappedBases=no ...  379     3e-103
  tig00000430 len=199451 reads=579 covStat=577.58 gappedBases=no ...  372     5e-101
  tig00000766 len=22030 reads=19 covStat=73.36 gappedBases=no cla...  365     8e-99
  tig00000820 len=161630 reads=606 covStat=354.09 gappedBases=no ...  360     3e-97
  tig00001382 len=143900 reads=490 covStat=362.02 gappedBases=no ...  351     2e-94
  tig00001054 len=130261 reads=335 covStat=377.80 gappedBases=no ...  345     7e-93
  tig00011987 len=22865 reads=63 covStat=54.78 gappedBases=no cla...  342     9e-92
  tig00000016 len=597014 reads=1702 covStat=1762.38 gappedBases=n...  338     1e-90
  tig00004245 len=46983 reads=101 covStat=145.13 gappedBases=no c...  322     8e-86
  tig00001199 len=110171 reads=303 covStat=326.23 gappedBases=no ...  322     8e-86
```

| | | |
|---|---|---|
| tig00004246 len=42866 reads=119 covStat=114.19 gappedBases=no c... | 320 | 3e-85 |
| tig00003701 len=66606 reads=166 covStat=197.05 gappedBases=no c... | 320 | 3e-85 |
| tig00001391 len=103712 reads=325 covStat=264.70 gappedBases=no ... | 318 | 1e-84 |
| tig00001318 len=116029 reads=80 covStat=485.40 gappedBases=no c... | 311 | 1e-82 |
| tig00017800 len=108736 reads=285 covStat=324.19 gappedBases=no ... | 306 | 6e-81 |
| tig00006142 len=24667 reads=24 covStat=94.12 gappedBases=no cla... | 306 | 6e-81 |
| tig00000980 len=120716 reads=595 covStat=176.64 gappedBases=no ... | 300 | 3e-79 |
| tig00001965 len=98243 reads=245 covStat=294.71 gappedBases=no c... | 295 | 1e-77 |
| tig00017693 len=414075 reads=1334 covStat=1097.73 gappedBases=n... | 282 | 7e-74 |
| tig00000963 len=116530 reads=374 covStat=300.03 gappedBases=no ... | 264 | 2e-68 |
| tig00004353 len=40397 reads=149 covStat=90.08 gappedBases=no cl... | 257 | 3e-66 |
| tig00000558 len=157957 reads=544 covStat=390.80 gappedBases=no ... | 253 | 3e-65 |
| tig00011500 len=28836 reads=45 covStat=104.73 gappedBases=no cl... | 250 | 4e-64 |
| tig00009898 len=14733 reads=6 covStat=39.68 gappedBases=no clas... | 239 | 8e-61 |
| tig00017737 len=159269 reads=462 covStat=435.38 gappedBases=no ... | 233 | 3e-59 |
| tig00019307 len=14647 reads=7 covStat=45.48 gappedBases=no clas... | 232 | 1e-58 |
| tig00001527 len=88379 reads=175 covStat=303.84 gappedBases=no c... | 232 | 1e-58 |
| tig00000093 len=354975 reads=783 covStat=1197.84 gappedBases=no... | 215 | 9e-54 |
| tig00017661 len=840329 reads=2776 covStat=2208.81 gappedBases=n... | 212 | 1e-52 |
| tig00004894 len=34499 reads=126 covStat=72.71 gappedBases=no cl... | 210 | 4e-52 |
| tig00006985 len=12414 reads=12 covStat=38.28 gappedBases=no cla... | 206 | 5e-51 |
| tig00000438 len=205342 reads=505 covStat=650.81 gappedBases=no ... | 201 | 2e-49 |

**>TRINITY_DN40142_c1_g1_i5**

CCTCTTAAAACGGAGCGTCCTCGCTATGCTGCTGCTACGAGTTGCTGTTGAAGACGGTTTATTTCGATTTCGTATAATTCGTTAATTTGTCATT
GTCTGTGAATCATTCTTCTATTTTCATAGTATCGGCAGAGATGTACCATACCATAAGTGATAACAACTAAAGTAAGGAAACCCATTATGATAAA
GAACGCAAAATTTACTGGGTTTTGTTCAATAGGTATCAAAAATGTTGACAGTTTTTGGATATTGCTGAATTTGTAGATTGAGTTTGAGGAAAGG
ATGCGGAGTACTTCAAGTTGTTCGCTATTTTAATTATGTAAGATTTCTCTGAGTTGTCGTTTATGATTGTAATATGTGATATTATCGATGGTTG
TAGATTCATTAAAGTGTATTAATTTAGGGCCTGATATATGTATGTTGTTCCAAATGTGCTGGTAATCTGTAAAAATGT

**BLASTn in PacBio Canu assembly:**

**Query=** TRINITY_DN40142_c1_g1_i5 Length=454

| Sequences producing significant alignments: | Score (Bits) | E Value |
|---|---|---|
| tig00011608 len=21095 reads=26 covStat=54.53 gappedBases=no cla... | 778 | 0.0 |
| tig00012318 len=15469 reads=14 covStat=42.62 gappedBases=no cla... | 769 | 0.0 |
| tig00005757 len=20510 reads=11 covStat=67.76 gappedBases=no cla... | 765 | 0.0 |
| tig00008422 len=11184 reads=7 covStat=36.15 gappedBases=no clas... | 738 | 0.0 |
| tig00001176 len=33509 reads=37 covStat=121.07 gappedBases=no cl... | 729 | 0.0 |
| tig00004871 len=21320 reads=15 covStat=74.63 gappedBases=no cla... | 720 | 0.0 |
| tig00013215 len=14752 reads=5 covStat=61.88 gappedBases=no clas... | 672 | 0.0 |
| tig00019780 len=21924 reads=41 covStat=53.03 gappedBases=no cla... | 655 | 0.0 |
| tig00004464 len=41343 reads=113 covStat=111.70 gappedBases=no c... | 655 | 0.0 |
| tig00010721 len=17700 reads=9 covStat=47.18 gappedBases=no clas... | 612 | 8e-174 |
| tig00008694 len=24345 reads=30 covStat=73.57 gappedBases=no cla... | 295 | 3e-78 |
| tig00000971 len=154460 reads=500 covStat=405.38 gappedBases=no ... | 251 | 3e-65 |
| tig00019422 len=18028 reads=7 covStat=50.51 gappedBases=no clas... | 212 | 3e-53 |
| tig00004439 len=44352 reads=79 covStat=154.29 gappedBases=no cl... | 185 | 4e-45 |

```
  tig00004978 len=48967 reads=78 covStat=166.03 gappedBases=no cl...  172  2e-41
  tig00013534 len=21574 reads=10 covStat=75.97 gappedBases=no cla...  154  6e-36
  tig00005959 len=35412 reads=35 covStat=129.01 gappedBases=no cl...  141  4e-32
  tig00002169 len=86941 reads=112 covStat=325.22 gappedBases=no c...  141  4e-32
  tig00019425 len=63695 reads=62 covStat=233.53 gappedBases=no cl...  125  3e-27
  tig00019156 len=39344 reads=65 covStat=130.28 gappedBases=no cl...  123  1e-26
  tig00018515 len=78919 reads=113 covStat=287.08 gappedBases=no c...  51.8 5e-05
```

The identified *MoY gene* (TRINITY_DN40292_c0_g3_i1 len=681) shows by BLASTn a highly similar sequence present in only one Canu assembly *FAM18* contig. See below:

**>TRINITY_DN40292_c0_g3_i1**

```
CGCTTAATATGTGCGATGTGTTATCACAGCCACGTTCAAGGCATTAACGCATTGCTTATTAAAAAACTTTATATTGTTTCGAGTACTGCTGATC
AGTATTATTATCCTGTGATTTAAGCGTTTATTTAAATAAAATTTGACATGGATATTGGAAATATTTCATCGAAAAATACAATCAGTTTAATAAC
AATAAAATATAACTCCAGAACTATCAAAGTAATTACTTCTAAAAGTCGTGGAATGGAACCGAAATTTTGGGGCAAAATGGAAATTGCAATGACA
GAAAATTATTTCGTAGAAGAAAAACCTCTTGTATACAATTTCGCAATAGAATATCGGAAATTAATGTCATAAATTTTGTGCAAGTCTGTTCACC
AAACATTCCTTTCATCCACATAATAACTCCGAAGGCATGCTGATACATTACAAAACAGAGTCAGAATATGATGAAACTCTTGGCTACATAACGG
AACACATGCTAGCAGATTGACCGGTAGTAGCTGTGGAAAAATATAAGGCATACCTAGTTTACTTAGTATTTTTTAAACTAAAAAACTTTTTGAA
TAAAATAATATAATAAGATACGATAATTTAGGAGCATTTTTAATAAATATAGTGGAACAAACAAGGTTATGTGTGACATGGAATTAACAAATTT
CGAAACTACTTTTGCTAAAGGGC
```

**BLASTn in PacBio Canu assembly:**

```
Query= TRINITY_DN40292_c0_g3_i1 Length=681
                                                                    Score      E
Sequences producing significant alignments:                        (Bits)   Value

  tig00013010 len=12420 reads=10 covStat=26.18 gappedBases=no cla...  1101    0.0
  tig00009898 len=14733 reads=6 covStat=39.68 gappedBases=no clas...  396     2e-108
  tig00019373 len=35768 reads=21 covStat=127.90 gappedBases=no cl...  244     7e-63
  tig00011001 len=12237 reads=39 covStat=22.27 gappedBases=no cla...  235     4e-60
  tig00019374 len=7827 reads=37 covStat=6.07 gappedBases=no class...  223     2e-56
  tig00010055 len=12860 reads=18 covStat=39.45 gappedBases=no cla...  221     8e-56
  tig00020818 len=11433 reads=10 covStat=25.59 gappedBases=no cla...  219     3e-55
  tig00019817 len=10935 reads=41 covStat=18.09 gappedBases=no cla...  219     3e-55
  tig00021013 len=6234 reads=8 covStat=15.05 gappedBases=no class...  215     3e-54
  tig00020139 len=27776 reads=73 covStat=68.73 gappedBases=no cla...  215     3e-54
```

This TRINITY_DN40292_c0_g3_i1 transcript used in a BLASTn on *Bactrocera oleae* XX and XY embryonic sexed transcriptomes, led to finding 4 weakly related transcripts from the same putative gene only in XY:

**BLASTn of TRINITY_DN40292_c0_g3_i1 on *B.oleae* XY transcriptome:**

```
> Query1 on c18307_g1_i5 len=473 path=[2135:0-71 4788:72-87 4835:88-158 2293:159-206
@4889@!:207-319 3350:320-321 @40@!:322-444 3879:445-449
3478:450-472]
Length=473

 Score = 41.0 bits (44),  Expect = 0.018
 Identities = 39/50 (78%), Gaps = 0/50 (0%)
 Strand=Plus/Plus

Query  182  TAATAACAATAAAATATAACTCCAGAACTATCAAAGTAATTACTTCTAAA  231
            |||||| |||||||||||| || |||||| | |   |||  |||||| ||
Sbjct  136  TAATAATAATAAAATATAATTCAAGAACTGTTATTATAACGACTTCTGAA  185

> Query1 on c18307_g1_i3 len=659 path=[2135:0-71 4788:72-87 4140:88-121 336:122-273
4835:274-344 2293:345-392 @4889@!:393-505 3350:506-507
@40@!:508-630 3879:631-635 3478:636-658]
Length=659

 Score = 41.0 bits (44),  Expect = 0.018
 Identities = 39/50 (78%), Gaps = 0/50 (0%)
 Strand=Plus/Plus

Query  182  TAATAACAATAAAATATAACTCCAGAACTATCAAAGTAATTACTTCTAAA  231
            |||||| |||||||||||| || |||||| | |   |||  |||||| ||
Sbjct  322  TAATAATAATAAAATATAATTCAAGAACTGTTATTATAACGACTTCTGAA  371

> Query1 on c18307_g1_i2 len=1599 path=[2135:0-71 4788:72-87 4140:88-121
336:122-273 4835:274-344 2293:345-392 @4889@!:393-505 2453:506-1302
3250:1303-1352 4432:1353-1407 3313:1408-1445 3350:1446-1447
@40@!:1448-1570 3879:1571-1575 3478:1576-1598]
Length=1599

 Score = 41.0 bits (44),  Expect = 0.018
 Identities = 39/50 (78%), Gaps = 0/50 (0%)
 Strand=Plus/Plus

Query  182  TAATAACAATAAAATATAACTCCAGAACTATCAAAGTAATTACTTCTAAA  231
            |||||| |||||||||||| || |||||| | |   |||  |||||| ||
Sbjct  322  TAATAATAATAAAATATAATTCAAGAACTGTTATTATAACGACTTCTGAA  371

> Query1 on c18307_g1_i1 len=789 path=[3683:0-24 @4661@!:25-161 2117:162-179
2135:180-251 336:252-403 4835:404-474 2293:475-522 @4889@!:523-635
3350:636-637 @40@!:638-760 3879:761-765 3478:766-788]
Length=789

 Score = 41.0 bits (44),  Expect = 0.018
 Identities = 39/50 (78%), Gaps = 0/50 (0%)
 Strand=Plus/Plus

Query  182  TAATAACAATAAAATATAACTCCAGAACTATCAAAGTAATTACTTCTAAA  231
            |||||| |||||||||||| || |||||| | |   |||  |||||| ||
Sbjct  452  TAATAATAATAAAATATAATTCAAGAACTGTTATTATAACGACTTCTGAA  501
```

A BLASTn of these 4 *B. oleae* overlapping transcripts sequences failed to find any corresponding transcripts in the XX *Bo* XX transcriptome, suggesting a very interesting XY-specificity of the corresponding gene expression.

BLASTn search of the PacBio Canu assembly, using medfly *MoY* 681 nt-long transcript sequence (including 5' and 3' UTRs) as DNA query (MK165756), identified a highly similar corresponding genomic sequence (95% identity). The *MoY* putative coding region of the transcript is 99% identical in the PacBio genomic sequence, with only 2 SNPs, with the second inducing a conservative amino acid substitution at position 63 (I->M) (Genbank acc. num. MK165755).

A BLASTn search using TRINITY_DN40292_c0_g3_i1 (derived from *Benakeion* strain) sequence on the male genome of FAM18 strain (PacBio Canu assembly of long reads) led to finding a unique 12 Kb long contig (tig00013010, len=12420), containing the whole *MoY* transcriptional unit, showing no introns and some polymorphism (95% DNA sequence identity; data not shown).

```
Query= TRINITY_DN40292_c0_g3_i1 Length=681
                                                                      Score      E
Sequences producing significant alignments:                          (Bits)  Value

  tig00013010 len=12420 reads=10 covStat=26.18 gappedBases=no cla...  1101 0.0
  tig00009898 len=14733 reads=6 covStat=39.68 gappedBases=no clas...  396  2e-108
  tig00019373 len=35768 reads=21 covStat=127.90 gappedBases=no cl...  244  7e-63
  tig00011001 len=12237 reads=39 covStat=22.27 gappedBases=no cla...  235  4e-60
  tig00019374 len=7827 reads=37 covStat=6.07 gappedBases=no class...  223  2e-56
  tig00010055 len=12860 reads=18 covStat=39.45 gappedBases=no cla...  221  8e-56
  tig00020818 len=11433 reads=10 covStat=25.59 gappedBases=no cla...  219  3e-55
  tig00019817 len=10935 reads=41 covStat=18.09 gappedBases=no cla...  219  3e-55
  tig00021013 len=6234 reads=8 covStat=15.05 gappedBases=no class...  215  3e-54
  tig00020139 len=27776 reads=73 covStat=68.73 gappedBases=no cla...  215  3e-54
  tig00019818 len=10394 reads=2 covStat=16.61 gappedBases=no clas...  215  3e-54
  tig00019375 len=16546 reads=64 covStat=23.92 gappedBases=no cla...  215  3e-54
  tig00019372 len=8510 reads=39 covStat=8.13 gappedBases=no class...  210  1e-52
  tig00014668 len=5823 reads=4 covStat=10.26 gappedBases=no class...  210  1e-52
  tig00011793 len=16988 reads=186 covStat=-59.19 gappedBases=no c...  210  1e-52
  tig00007607 len=12605 reads=4 covStat=37.89 gappedBases=no clas...  210  1e-52
  tig00020587 len=14310 reads=42 covStat=35.74 gappedBases=no cla...  159  2e-37
  tig00010611 len=8603 reads=2 covStat=13.53 gappedBases=no class...  95.1 8e-18
  tig00003532 len=50280 reads=147 covStat=134.34 gappedBases=no c...  53.6 2e-05
```

The BLASTn found also 4 shorter *MoY* weakly related sequences with the 12 Kb long contig, showing 70% identity over 200-500 bp long regions (the 4 sequences are at positions 5 Kb, 8 Kb, 9 Kb and 11 Kb in the 12 Kb long contig; see the 4 dashed red lines in Fig. 1c). The whole *MoY* transcribed region (0.7 Kb) seems to be a single copy in the Canu assembly and absent in the available medfly genome at NCBI. The *MoY* putative coding region of the transcript (*Benakeion* strain) is 99% identical in the FAM18 male genomic sequence, with only 2 SNPs, with the second inducing a conservative amino acid substitution at position 63 (I->M).

A tBLASTn search with MOY amino acid sequence in the NCBI *Ceratitis* reference genome or transcriptome (refseq_RNA) failed to find respectively *MoY* gene or RNA identical copies, but only an unplaced genomic scaffold (NW_019377179.1) containing a 1 Kb long DNA sequence which potentially encodes for a very short MOY related protein sequence (79% aa identity over a 19 aa long region).

A BLASTn search with the *MoY* Trinity contig sequence in the FAM18 assembly found a genomic contig (tig00009898 len=14733 bp) containing only a truncated (230 bp long) but identical *MoY* sequence and a nearby duplication of similar length, showing 70% sequence identity. The 200 bp 5' region of *MoY* (5'UTR and first 20 amino acid coding region ) has related sequences (70-80% identity) in 15 other Canu genomic contigs, showing often multiple copies with the same contig, suggesting that translocations and duplications of *MoY* truncated versions occurred.

A tBLASTn search with MOY amino acid sequence in the PacBio Canu assembly led to finding the previous 12 Kb long contig, as expected, but also other 18 contigs (6-35 Kb long) containing putative MOY-related shorter ORFs which correspond only to truncated versions (20-40 aa long, showing 50-70% aa identity), as expected on the basis of the previous BLASTn analysis.

BLASTn search showed that the *MoY* gene has an identical unique sense transcript (see below for ORF) which is 681 bp long in the mixed XX/XY 4-6 h embryonic transcriptome, with 2 shorter antisense RNAs (see Fig. 1b; stranded RNA sequencing made possible to identify the antisense RNAs), overlapping/pairing with *MoY* respectively in the 5' UTR and 3' UTR. No *MoY* transcripts have been found in XX-only embryonic *Ceratitis* transcriptome. One of the potential ORFs present in the *MoY* Trinity contig corresponds to a putative 70 aa long protein, which was later found conserved in other Tephritidae species (see Supplementary Information Notes 5-12).

```
>MOY_70_aa
MDIGNISSKNTISLITIKYNSRTIKVITSKSRGMEPKFWGKMEIAMTENYFVEEKPLVYNFAIEYRKLMS
```

A BLASTn analysis of TRINITY_DN40292_c0_g3_i1 to search the first 7 M factor candidates (Extended Data Table 1), led to finding their identity with the last transcript of the list (*corvus*) (Supplementary Information Note 1), which however escaped our attention and interest, at the time of the first analysis.

**>corvus-A (comp172828_c0_seq1 len=411)**

TATATTTTTCCACAGCTACTACCGGTCAATCTGCTAGCATGTGTTCCGTTATGTAGCCAAGAGTTTCATCATATTCTGAC
TCTGTTTTGTAATGTATCAGTATGCCTTCGGAGTTATTATGTGGCAGATATATTTGAAAGGAATGTTTGGTGAAAAGACT
TGCACAAAATTTATGACATTAATTTCCGATATTCCATTGCGAAATTGTATACAAGAGGTTTTTCTTCTACGAAATAATTT
TCTGTCATTGCAATTTCCATTTTGCCCCAAAATTTCGGTTCCATTCCACGACTTTTAGAAGTAATTACTTTGATAGTTCT
GGAGTTATATTTTATTGTTATTAAACTGATTGTATTTTTCGATGAAATATTTCCAATATCCATGTCAAATTTTATTTAAA
TAAACGCTTAA

**corvus-A (comp172828_c0_seq1 len=411)**

**Sequence ID:** Query_73779 **Length:** 411 **Number of Matches:** 1

| Score | Expect | Identities | Gaps | Strand |
|---|---|---|---|---|
| 684 bits(370) | 0.0 | 399/411(97%) | 9/411(2%) | Plus/Minus |

```
Query  114   TTAAGCGTTTATTTAAATAAAATTTGACATGGATATTGGAAATATTTCATCGAAAAATAC  173
             ||||||||||||||||||||||||||||||||||||||||||||||||||||||||||||
Sbjct  411   TTAAGCGTTTATTTAAATAAAATTTGACATGGATATTGGAAATATTTCATCGAAAAATAC  352

Query  174   AATCAGTTTAATAACAATAAAATATAACTCCAGAACTATCAAAGTAATTACTTCTAAAAG  233
             ||||||||||||||||||||||||||||||||||||||||||||||||||||||||||||
Sbjct  351   AATCAGTTTAATAACAATAAAATATAACTCCAGAACTATCAAAGTAATTACTTCTAAAAG  292

Query  234   TCGTGGAATGGAACCGAAATTTTGGGGCAAAATGGAAATTGCAATGACAGAAAATTATTT  293
             ||||||||||||||||||||||||||||||||||||||||||||||||||||||||||||
Sbjct  291   TCGTGGAATGGAACCGAAATTTTGGGGCAAAATGGAAATTGCAATGACAGAAAATTATTT  232

Query  294   CGTAGAAGAAAAACCTCTTGTATACAATTTCGCAATAGAATATCGGAAATTAATGTCATA  353
             |||||||||||||||||||||||||||||||||||| |||||||||||||||||||||||
Sbjct  231   CGTAGAAGAAAAACCTCTTGTATACAATTTCGCAATGGAATATCGGAAATTAATGTCATA  172

Query  354   AATTTTGTGCAAGTCTGTTCACCAAACATTCCTTTCA--T-------CCACATAATAACT  404
             |||||||||||||||| ||||||||||||||||||||   |       |||||||||||||
Sbjct  171   AATTTTGTGCAAGTCTTTTCACCAAACATTCCTTTCAAATATATCTGCCACATAATAACT  112

Query  405   CCGAAGGCATGCTGATACATTACAAAACAGAGTCAGAATATGATGAAACTCTTGGCTACA  464
                |||||||||| |||||||||||||||||||||||||||||||||||||||||||||||||
Sbjct  111   CCGAAGGCATACTGATACATTACAAAACAGAGTCAGAATATGATGAAACTCTTGGCTACA  52

Query  465   TAACGGAACACATGCTAGCAGATTGACCGGTAGTAGCTGTGGAAAAATATA  515
             |||||||||||||||||||||||||||||||||||||||||||||||||||
Sbjct  51    TAACGGAACACATGCTAGCAGATTGACCGGTAGTAGCTGTGGAAAAATATA  1
```

Also, a BLASTn search with the *MoY* Trinity sequence in the 4-6 h XX/XY mixed embryonic transcriptome led to identifying 8 different RNA contigs (0.2-2.8 Kb long), showing 70-87% identity over 150-300 nt long regions and having either sense or antisense orientation. 6 out of 8 contigs showed DNA similarity in regions containing the *MoY* ORF region. A tBLASTn search with MOY amino acid sequence in the mixed XX/XY embryonic transcriptome led to finding again the same 6 RNA contigs, encoding for truncated and divergent MOY sequences. In contrast, *MoY* BLASTn and MOY tBLAST searches in the XX-only embryonic transcriptome failed to identify RNA contigs with significant similarity (DNA regions >100 bp; data not shown). These observations suggested that the 8 RNA contigs, weakly related to *MoY*, could correspond to novel medfly Y-linked transcribed sequences.

**Supplementary Note 2 | A preliminary screening of M candidates and list of data available at NCBI used for *Ceratitis capitata* embryonic (0-48 h AEL) and adult male transcriptome assembly (ISPRA strain).**

To identify the medfly *M* factor, an approach similar to the chromosome quotient (CQ) was utilized. Chromosome quotients were calculated for all transcripts *de novo* assembled from embryonic (0-48 h old) and adult male RNA-seq data (available at NCBI). Chromosome quotients were calculated using the methods described in Hall et al., (2013). Briefly, Illumina *Fam18* genomic reads, separately sequenced from male and females, were aligned to each transcript arising from *de novo* assembly with high stringency using bowtie1 with -v 0 flag. Then, the ratio of female to male alignments was calculated for each transcript. A transcript was considered likely to have arisen from the Y chromosome if it had 30 or more alignments from male sequencing data and less than 3 alignments from female sequencing data. This initial attempt to identify the medfly *M* factor relied on limited RNA-seq (no biological replicates and 0-48 h old embryos) and male/female genomic data available at NCBI and led to the identification of 7 Y-linked male-specific transcriptional units, most of which likely corresponded to pseudogenes, with 4 confirmed to be Y-linked by PCR on gDNA (data not shown). Most of these are absent in the available medfly assembled genome, as expected for medfly Y-linked genes, considering the technical difficulties to assemble Y-derived sequences from repetitive regions also observed in other species. Moreover, in 5 of these transcripts, BLASTn analysis showed similarity (70-90%) to duplicated paralogous sequences present in the 0-48 h assembled embryonic transcriptome (Extended Data Table 1). BLASTx analyses of these 5 genes on *C. capitata* and *D. melanogaster* protein databases showed that they seem to correspond to transcribed pseudogenes, having only short stretches of similarity to known proteins, while other 2 (*corvus* and *dorado*) showed no protein similarity neither multiple copies.

Following a list of data available at NCBI used for the assembly of a transcriptome from *C. capitata* embryos 0-48 h after egg laying (AEL) and adult male RNA-seq data (both from ISPRA strain; Pavia, Italy: SRX272876; SRX272878). List of *Ceratitis capitata* male genomic data available at NCBI and used in this study: SRX276046; SRX275788; SRX272878. List of *C. capitata* female genomic data available at NCBI and used in this study: SRX275787; SRX276048; SRX276047.

A table listing the 7 embryonic 0-48 h AEL/male adult transcripts corresponding to putative Y-linked genes, selected by CQ-like analysis, is reported below. Their presence/absence in the available medfly genome were analysed by BLASTn. Presence of paralogous transcripts in the 0-48 h AEL embryonic transcriptome was analysed by BLASTn. The presence of conserved putative ORFs was analysed by BLASTx (BLOSUM45) in *C. capitata* and *D. melanogaster* protein databases (considering only the hits with E value < 2.6).

| CQ selected transcripts 0-48 h embryos+males | Medfly Baylor Genome | Multiple paralogous contigs in 0-48 embryonic transcriptome | BLASTx Ceratitis (E value < 2.6) | BLASTx Drosophila (E value < 2.6) |
|---|---|---|---|---|
| Orion | no | 18 contigs (at least 7e-59) | cytosol aminopeptidase-like XP_023159293.1 | Sperm-Leucylaminopeptidase 3, isoform C |
| | | | myb-like protein I | (E value 1e-13) |
| | | | XP_004522636.1 (E value 1e-16) | NP_648394.1 |
| | | | zinc finger protein 239 (E value 2e-4) | |
| Lyra | yes | 6 contigs (at least 2e-44) | putative gustatory receptor 59f | none |
| | | | XP_004526066.1 (E value 2.5) | |
| Aries (Orion B related) | no | 4 contigs (at least 4e-63) | cytosol aminopeptidase-like XP_023159293.1 (E value 5e-36) | Sperm-Leucylaminopeptidase 3, isoform C |
| | | | | NP_648394.1 (E value 2e-15) |
| Dorado | yes | none | none | none |
| Pavo (94% identical to Orion B) | no | 4 contigs (2e-49) | cytosol aminopeptidase-like XP_023159293.1 (8e-06) | Sperm-Leucylaminopeptidase 3, isoform C |
| | | | | NP_648394.1 (E value 0.62) |
| Norma | no | 1 contig (1e-82) | NADH dehydrogenase (ubiquinone) chain 1 (mitochondrion) | NADH dehydrogenase subunit 1 (mitochondrion) |
| | | | CAB45100.1 (E value 4e-15) | Sequence ID: YP_009047278.1 (E value 7e-14) |
| Corvus | no | none | none | none |

**Supplementary Note 3 | DNA polymorphism of *MoY* sequences from *Benakeion* and *Fam18* strains.**

The *MoY* AUG and STOP codons are in bold. The *MoY* putative coding region of the transcript is 99% identical, with only 2 SNPs, with the second inducing a conservative amino acid substitution at position 63 (I->M).

```
BEN-MOY   1    CGCTTAATATGTGCGATGTGTTATCACAGCCACGTTCAAGGCATTAACGCATTGCTTATT  60
               |||||||||||||||||||||||||||||||||||||||| ||||||||||||||| |||
FAM18MOY 190   CGCTTAATATGTGCGATGTGTTATCACAGCCACGTTCAAGACATTAACGCATTGCTCATT  249

BEN-MOY   61   AAAAAACTTTATATTGTTTCGAGTACTGCTGATCAGTATTATTATCCTGTGATTTAAGCG  120
               |||||||||||| ||||||||||||||||||||||||||||| ||||||||||||||||||
FAM18MOY 250   AAAAAACTTTATCTTGTTTCGAGTACTGCTGATCAGTATTACTATCCTGTGATTTAAGCG  309

BEN-MOY   121  TTTATTTAAATAAAATTTGACATGGATATTGGAAATATTTCATCGAAAAATACAATCAGT  180
               |||||||||||||||||||||||||||||||||||||||||||||||||| |||||||||
FAM18MOY 310   TTTATTTAAATAAAATTTGACATGGATATTGGAAATATTTCATCGAAAAACACAATCAGT  369

BEN-MOY   181  TTAATAACAATAAAATATAACTCCAGAACTATCAAAGTAATTACTTCTAAAAGTCGTGGA  240
               ||||||||||||||||||||||||||||||||||||||||||||||||||||||||||||
FAM18MOY 370   TTAATAACAATAAAATATAACTCCAGAACTATCAAAGTAATTACTTCTAAAAGTCGTGGA  429

BEN-MOY   241  ATGGAACCGAAATTTTGGGGCAAAATGGAAATTGCAATGACAGAAAATTATTTCGTAGAA  300
               ||||||||||||||||||||||||||||||||||||||||||||||||||||||||||||
FAM18MOY 430   ATGGAACCGAAATTTTGGGGCAAAATGGAAATTGCAATGACAGAAAATTATTTCGTAGAA  489

BEN-MOY   301  GAAAAACCTCTTGTATACAATTTCGCAATAGAATATCGGAAATTAATGTCATAAATTTTG  360
               ||||||||||||||||||||||||||||| ||||||||||||||||||||||||||||||
FAM18MOY 490   GAAAAACCTCTTGTATACAATTTCGCAATGGAATATCGGAAATTAATGTCATAAATTTTG  549

BEN-MOY   361  TGCAAGTCTGTTCACCAAACATTCCTTTCA--T-------CCACATAATAACTCCGAAGG  411
               ||||||||| ||||||||||||||||||||  |       ||||||||||||||||||||
FAM18MOY 550   TGCAAGTCTTTTCACCAAACATTCCTTTCAAATATATCTGCCACATAATAACTCCGAAGG  609

BEN-MOY   412  CATGCTGATACATTACAAAACAGAGTCAGAATATGATGAAACTCTTGGCTACATAACGGA  471
               ||| ||||||||||||||||||||||||||||||||||||||||||||||||||||||||
FAM18MOY 610   CATACTGATACATTACAAAACAGAGTCAGAATATGATGAAACTCTTGGCTACATAACGGA  669

BEN-MOY   472  ACACATGCTAGCAGATTGACCGGTAGTAGCTGTGGAAAAATATAAGGCATACCTAGTTTA  531
               |||||||||||||||||||  ||||||||||   ||||||||||||||||||||||||||
FAM18MOY 670   ACACATGCTAGCAGATTGATTGGTAGTAGCT---GAAAAATATAAGGCATACCTAGTTTA  726

BEN-MOY   532  CTTAGTATTTTTTAAACT---aaaaaactttttgaataaaataatataataagatacgat  588
               |||||| |||||||||||   |||||||||||| ||||||||||||||||||||||||||
FAM18MOY  27   CTTAGTCTTTTTTAAACTAAAAAAAAACTTTTTAAATAAAATAATATAATAAGATACGAT  786

BEN-MOY   589  aatttaggagcatttttaataaatataGTGGAACAAACAAGGTTATGTGTGACATGGAAT  648
               ||||||||||||||||||||||||||||||||||||||||||||||||||||||||||| 
FAM18MOY 787   TATTTAGGAGCATTTTTAATAAATATAGTGGAACAAACAAGGTTATGTGTGACATGGAAG  846

BEN-MOY   649  TAACAAATTTCGAAACTACTTTTGCTAAAGGGC  681
               | |||||| |||||||||||||||||||| |||
FAM18MOY 847   TGACAAATATCGAAACTACTTTTGCTAAAAGGC  879
```

**Supplementary Note 4 | Other novel transcriptional units in the *MoY* genomic flanking regions.**

A CQ analysis, mapping male and female DNA Illumina reads on the 12 Kb long genomic contig (tig00013010) (data not shown), as well as PCRs on male and female genomic DNA, confirmed that this contig seems to be derived from the Y chromosome (data not shown). Furthermore, XX/XY embryos but not XX-only embryos Illumina reads map along the 12 Kb contig, indicating the presence of novel male-specific RNAs produced from this region (data not shown; see below for Illumina transcripts sequences). Indeed, a BLASTn of the whole 12 Kb region on the available NCBI medfly genome and on NCBI related RNA reference database failed to find sequence similarity confirming the novelty of the identified genomic and related RNA sequence information.

The 12 Kb long Canu genomic sequence was used in a BLASTn analysis on *Ceratitis* mixed XX/XY embryonic transcriptome. 20 Trinity transcripts belonging to 10 different genes, have been mapped along the genomic region (List in the table below). Two genes (DN40516 in green and DN40292 in violet) have 5 duplicated copies of variable length along the region, and they overlap with various extent in sense-antisense orientation (violet and green RNAs; Fig. 1b). The relative positions of the 10 RNAs listed above along the 12 Kb long Y-specific genomic region are indicated in Fig. 1b. A colour code of the RNAs in this list and of the arrows representing them in Fig. 1a can help to localize them.

| RNA | Name | Lenght | Position | BLASTn female embryos | BLASTn NCBI hits | BLASTx NCBI hits | BLASTn Bo male embryos hits | BLASTn Bo female embyos hits |
|---|---|---|---|---|---|---|---|---|
| 1 | TRINITY_DN26767_c0_g1_i1 | 381 bp | 84-381 | none at 100% | yes | yes (homeobox) | **yes** | **yes** |
| 2 | TRINITY_DN45758_c0_g1_i1 | 212 bp | 1119-1330 | none at 100% | none | yes (phosphiolipase) | none | none |
| 3 | TRINITY_DN6507_c0_g1_i1 | 244 bp | 1444-1687 | none at 100% | none | none | none | none |
| 4 | TRINITY_DN40292_c0_g1_i9 | 2865 bp | 4837-6083<br>6098-7025<br>8721-9610<br>9609-11390<br>12145-12420 | none at 100% | yes | none | none | none |
| 5 | TRINITY_DN40516_c0_g1_i5 | 1990 bp | 4443-6083<br>5112-6267<br>8323-9610<br>9609-10244 | none | none | none | none | none |
| 6 | TRINITY_DN32944_c0_g1_i1 | 467 bp | 8291-8747<br>4443-4861 | none | none | none | none | none |
| 7 | TRINITY_DN38978_c0_g1_i1 | 535 bp | 6854-7338 | none at 100% | yes | | none | none |
| 8 | TRINITY_DN40292_c0_g3 | 681 bp | 3620-4309 | none | none | none | **yes** | none |
| 9 | TRINITY_DN77369_c0_g1_i1 | 243 bp | 3760-3518 | none | none | none | none | none |
| 10 | TRINITY_DN104942_c0_g1_i1 | 285 bp | 4263-3970 | none | none | none | none | none |

**Supplementary Note 5 | *MoY* orthologue in the olive fly *Bactrocera oleae* (*BoMoY*).**

Only three out of 13 Trinity transcripts showed sequence similarity to male but not female embryonic transcripts of the related Tephritidae species, *Bactrocera oleae* (Extended Data Table 2). First search by BLASTp and tBLASTn at NCBI protein and nt databases, using MOY/*MoY* sequences as probes, failed to find homologous or weakly similar sequences with some statistical significance. In contrast, tBLASTx search using MOY sequence in transcriptomes that we assembled from SRA databases and in WGS databases of 14 Tephritidae species led to identifying putative *MoY* orthologues in 8 of them (Fig. 3a). A BLASTn and tBLASTn search with *MoY* in the XY embryonic transcriptome of *Bactrocera oleae* (assembled from SRA SRX265053 downloaded from NCBI) led to finding *MoY* orthologous transcripts (*BoMoY*) and putative encoded protein (BoMOY, 87 aa), showing respectively 77% DNA identity over a 57 nt long region and 57% protein similarity. On the contrary, the sequences of 2 antisense *MoY* RNAs (corresponding to the *MoY* 5' and 3' UTRs) and the other transcripts present in the flanking *Ceratitis MoY* region are not conserved in the *B. oleae* sexed embryonic transcriptome. A BLASTp analysis showed that MOY and BoMOY shares 63% aa similarity over a 58 aa long region (see below).

```
Identities 21/58(36%)
Positives 37/58(63%)

MOY     11     TISLITIKYNSRTIKVITSKSRGMEPKFWGKMEIAMTENYFVEEKPLVYNFAIEYRKL  68
               ++ +I IKYNSRT+ + TS+ R M  + W   E   T+ + +++K +V N + E++KL
BoMOY   6      SVWIIIIKYNSRTVIITTSERRIMPRRVWNAKE---TKPH-IKKKQMVLNLSTEFKKL  59
```

Interestingly, *BoMoY* transcripts were found in the XY but not in the XX embryos transcriptomes (XX embryonic transcriptome assembled from SRA SRX265052 downloaded from NCBI), indicating possibly male-specific Y-linked conservation. *BoMoY* gene is partially contained within a 1 Kb long genomic scaffold found by BLASTn at NCBI WGS sequence (Sequence ID: JXPT01043932.1). Differently to *MoY*, *BoMoY* seems to be an intron-containing gene. A preliminary draft of the *BoMoY* gene suggests the presence of 2 introns (a first 186 nt long in the 5' UTR region and a second 941 nt long within the ORF region) and potentially encodes for a 87 aa long BoMOY protein. The longer transcript (1.6 Kb) seems to correspond to a BoMoY unspliced isoform, potentially encoding for a shorter protein isoform (BoMOY-2; 71 aa long). *BoMoY*-specific PCR on sexed *B. oleae* genomic DNA confirmed that the putative gene is Y-linked also in this other Tephritidae species (see

Fig. 3c). On the contrary, the sequences of 2 antisense *MoY* RNAs are not conserved in *B. oleae* sexed embryonic transcriptome, neither the sequences of other transcripts present in the flanking *Ceratitis MoY* genomic region.

A tBLASTn search with BoMOY longer putative amino acid sequence (87 aa long) in the NCBI *B. oleae* whole-genome shotgun led to finding 2 assembled genomic contigs, 1 kb and 7 Kb long, respectively (sequence ID: JXPT01043932.1 and LGAM01008500.1), containing only the 5’ and 3’ *BoMoY* regions. A third 6 Kb long assembled genomic contig (LGAM01009849.1), contains at one of its very end *BoMoY* fragment encoding 15 BoMOY aa sequence of the C-terminus. A similar tBLASTn search in the male XY *B. oleae* embryonic transcriptome (assembled from SRA at NCBI Accessions: SRX265052, female embryos, and SRX265053, male embryos), led to finding only a unique Trinity contig (with 5 Trinity 5 isoforms; see list of sequences: BoMoY Trinity isoforms). Hence, we have found no indications of duplicated and divergent *BoMoY* related sequences, differently to medfly *MoY*. PCR on sexed genomic DNA of the olive fly confirmed that *BoMoY* is Y-linked.

**BoMOY_87_aa**

MDKMRSVWIIIIKYNSRTVIITTSERRIMPRRVWNAKETKPHIKKKQMVLNLSTEFKKLKNKKCLFARKFSFLPFSQGNNCRLQHLQ

**BoMoY_short_71_aa**

MDKMRSVWIIIIKYNSRTVIITTSERRIMPRRVWNAKETKPHIKKKQMVLNLSTEFKKLKNKKCLFARKFR

**Supplementary Note 6 | *MoY* orthologue in *Bactrocera jarvisi*, a mango pest native to Australia.**

A tBLASTn analysis of a XY male and a XX female *Bactrocera jarvisi* 3-5 h embryonic transcriptomes (assembled from NCBI SRAs XY male embryos replicates: SRX697431 and SRX697428; and SRAs XX 3-5 h female embryos replicates: SRX697435 and SRX697434) led to finding only in the XY embryos a Trinity contig (including 3 Trinity isoforms; see List of RNAs) encoding for a BjMOY protein (70 aa) and showing by BLASTp an overall 76% aa similarity to BoMOY and 60% to MOY (see below).

```
Identities 44/69(64%)
Positives 53/69(76%)

BoMOY   4       MRSVWIIIIKYNSRTVIITTSERRIMPRRVWNAKETKPHIKKKQMVLNLSTEFKKL---K  60
                M SVWIII K+NSRTVI+ +SER IM R+ WN K  KP I++K+++LNLSTEFKKL
BjMOY   1       MGSVWIIIRKHNSRTVILASSERLIMSRKFWNEKNLKPDIEEKEIILNLSTEFKKLMNNN  60

BoMOY  61       NKKCLFARK  69
                N KCLF RK
BjMOY  61       NTKCLFTRK  69

Identities  19/60(32%)
Positives  36/60(60%)

MOY     11      TISLITIKYNSRTIKVITSKSRGMEPKFWGKMEIAMTENYFVEEKPLVYNFAIEYRKLMS  70
                ++ +I  K+NSRT+ + +S+   M  KFW +  +       +EEK ++ N + E++KLM+
BjMOY  3        SVWIIIRKHNSRTVILASSERLIMSRKFWNEKNLKPD----IEEKEIILNLSTEFKKLMN  58
```

The three Trinity isoforms contain a stop codon in third aa position, following the putative AUG; however, a search of the 3 SRA male embryos libraries led to finding SRAs containing a codon for serine in place of the stop codon (See List of *MoY* RNAs). Hence, we speculated that highly similar duplicated copies of *BjMoY* are present in the Bj genome, with some containing a stop codon in the third aa position of the putative ORF. Hence, Trinity assembly preferred to compose transcript containing those SRAs most numerically represented. It is expected that a male determining factor amplify horizontally to escape inactivation by mutations, as observed for example in *M. domestica* (Sharma et al., 2017). Hence considering the very high sequence identity and the existence of those mentioned SRAs, we speculated that a *BjMoY* transcriptional active copy containing a full length BjMOY protein is present in *B. jarvisi* genome. Hence, we manually replaced the stop codon with a serine in the BjMOY putative protein and considered it as a concrete reference. The 3 Trinity isoforms seem to be derived by alternative splicing involving intron/exon regions localized in *BjMoY* 3' UTR. No *B. jarvisi* genomic sequences are presently available at NCBI.

**BjMOY 70 aa**

MGSVWIIIRKHNSRTVILASSERLIMSRKFWNEKNLKPDIEEKEIILNLSTEFKKLMNNNNTKCLFTRKV

**Supplementary Note 7 | *MoY* orthologue in the Queensland fruit fly *Bactrocera tryoni*, native to Australia (Q-fly), which affects mostly pome, stone and citrus fruits.**

A tBLASTn with *BdMOY* aa sequence on NCBI WGS database of another Australian species, *Bactrocera tryoni*, (Qfly, Queensland fly) led to find a 7 Kb long genomic sequence (GenBank: JHQJ01009763.1) showing an overall 97% protein similarity. The *BtMoY* putative coding region seems to be entirely contained in the genomic contig and a highly similar second *BtMoY* coding region (94% nt identity; *BtMoY-2*) is also present at a distance of 3.5 Kb from the first and on the same putative transcription orientation coding for a shorter putative BtMOY-2 protein (55 aa long). Apparently, no other *BtMoY* copies are present in the WGS database of *Bactrocera tryoni,* when searched by BLASTn or tBLASTn.

*BtMoY* shares 85% nt sequence identity with *BjMoY,* of the other Australian species *B. jarvisi*, over a 700 nt long region. The 2 MOY proteins share an overall 94% aa sequence similarity.

```
Identities 61/69(88%)
Positives  65/69(94%)

BtMOY  1   MGSVLIIIRKHNSRTVILTSSERLIMSRRFWNEKNMKPDIEEKEMVLNLSTEFKKLMNNN  60
           MGSV IIIRKHNSRTVIL SSERLIMSR+FWNEKN+KPDIEEKE++LNLSTEFKKLMNNN
BjMOY  1   MGSVWIIIRKHNSRTVILASSERLIMSRKFWNEKNLKPDIEEKEIILNLSTEFKKLMNNN  60

BtMOY  61  NKKYLFTRK  69
           N K LFTRK
BjMOY  61  NTKCLFTRK  69
```

**BtMOY 70 aa**

```
MGSVLIIIRKHNSRTVILTSSERLIMSRRFWNEKNMKPDIEEKEMVLNLSTEFKKLMNNNNNKKYLFTRKF
```

**Supplementary Note 8 | *MoY* orthologue in the Oriental fruit fly *Bactrocera dorsalis* (it affects a broad range of host fruits; endemic of South East Asia; invasive in USA territories).**

A tBLASTn analysis of the oriental fly *Bactrocera dorsalis* SRA NCBI databases, using *BjMOY* as probe, led to find a SRA sequence, showing very high aa sequence identity (SRA: SRR316210.7953824.1 and SRA: SRR316210.7953824.2; SRX085118, *Bactrocera dorsalis* transcriptome analysis). PCR on male and female genomic DNA of *B. dorsalis*, confirmed that this sequence is Y-specific (Fig. 4a). The whole *BdMoY* ORF coding region was cloned by RT-PCR from embryonic *B. dorsalis* RNA, using 2 primers designed on the forward and reverse SRA sequence (SRA: SRR316210.7953824.1 and SRA: SRR316210.7953824.2; Supplementary Methods Table 1), identified as highly similar to *BjMOY* by tBLASTn (SRX085118; *Bactrocera dorsalis* transcriptome analysis. BLASTp analyses showed that BdMOY (70 aa) is similar to BjMOY (97% aa overall similarity), BoMOY (80% aa overall similarity) and MOY (60% aa similarity over a 55 aa long region) (see below).

**BdMOY 70 aa**

```
MGSVWIIIRKHNSRTVILTSSQRLLLSRRFWNEKNMKPDIEEKEIVLNLSTEFKKLMNNNNKKCLFTRKF

Identities 61/69(88%)
Positives  67/69(97%)

BdMOY  1       MGSVWIIIRKHNSRTVILTSSQRLLLSRRFWNEKNMKPDIEEKEIVLNLSTEFKKLMNNN  60
               MGSVWIIIRKHNSRTVIL SS+RL++SR+FWNEKN+KPDIEEKEI+LNLSTEFKKLMNNN
BjMOY  1       MGSVWIIIRKHNSRTVILASSERLIMSRKFWNEKNLKPDIEEKEIILNLSTEFKKLMNNN  60

BdMOY  61      NKKCLFTRK  69
               N KCLFTRK
BjMOY  61      NTKCLFTRK  69

Identities 46/70(66%)
Positives 56/70(80%)

BdMoY  1       MGSVWIIIRKHNSRTVILTSSQRLLLSRRFWNEKNMKPDIEEKEIVLNLSTEFKKLMNNN  60
                M SVWIII K+NSRTVI+T+S+R ++ RR WN K  KP I++K++VLNLSTEFKKL
BoMOY  4        MRSVWIIIIKYNSRTVIITTSERRIMPRRVWNAKETKPHIKKKQMVLNLSTEFKKL---K  60

BdMoY  61       NKKCLFTRKF  70
                NKKCLF RKF
BoMOY  61       NKKCLFARKF  70

Identities 18/60(30%)
Positives  36/60(60%)
BdMoY  3        SVWIIIRKHNSRTVILTSSQRLLLSRRFWNEKNMKPD----IEEKEIVLNLSTEFKKLMN  58
                ++ +I  K+NSRT+ + +S+   +  +FW +  +       +EEK +V N + E++KLM+
MOY     11      TISLITIKYNSRTIKVITSKSRGMEPKFWGKMEIAMTENYFVEEKPLVYNFAIEYRKLMS  70
```

**Supplementary Note 9 | *MoY* orthologue in the melon fly *Zeugodacus cucurbitae*, (ex *Bactrocera cucurbitae*), which is native of India and present in South-East Asia, as well as Japan, Australia, Hawaii.**

A tBLASTn with BdMOY aa sequence on NCBI WGS database of *Zeugodacus cucurbitae* led to find a 1.3 Kb long genomic sequence (GenBank: JRNW01040954.1) showing 66% aa similarity over 24 aa long amino terminus region and 84% over a 19 aa long central region. A frame shift is observed within the *ZcMoY* putative genomic region, possibly due to sequencing error, or to presence of a small intron, or existence of multiple *ZcMoY* copies with some inactivated by mutations. The two *ZcMoY* coding regions shifted by 1 open reading frame contain 30 aa long and 43 aa long ORFs, respectively. A BLASTp alignment of the 2 ZcMOY sequences with BdMOY led to join them into a 54 aa long putative ZcMOY, presuming DNA sequencing error in the genomic sequence. No corresponding *ZcMoY* transcript sequences were found in the few *Z. cucurbitae* SRA databases available at NCBI. BLASTp analysis showed that ZcMOY (54 aa) is similar to BdMOY (70% aa overall similarity) and to MOY (60% aa similarity over a 55 aa long region) (see below).

**ZcMoY 54**

```
MGSVWVLKTKYNSRTITVTTSERKPISSIFFWNAKNTHLHIETKHIVFNLTTEF

Identities 29/54(54%)
Positives 38/54(70%)
ZcMOY  1   MGSVWVLKTKYNSRTITVTTSERKPISSIFFWNAKNTHLHIETKHIVFNLTTEF  54
           MGSVW++  K+NSRT+ +T+S+R  +S   FWN K     IE K IV NL+TEF
BdMOY  1   MGSVWIIIRKHNSRTVILTSSQRLLLSR-RFWNEKNMKPDIEEKEIVLNLSTEF  53

18/56(32%)
30/56(53%)
ZcMOY  3   SVWVLKTKYNSRTITVTTSERKPISSIFFWN----AKNTHLHIETKHIVFNLTTEF  54
           ++ ++  KYNSRTI V TS+ + +  FW       A   +  +E K +V+N   E+
MOY    11  TISLITIKYNSRTIKVITSKSRGMEPK-FWGKMEIAMTENYFVEEKPLVYNFAIEY  65
```

**Supplementary Note 10 | *MoY* orthologue in *Bactrocera latifrons*, native of Asia, which is an invasive pest of fruit and vegetables, mainly belonging to Solanaceae, including tomato, and to a lesser extent to Cucurbitaceae.**

A tBLASTn with BdMOY aa sequence on NCBI WGS database of another Asian species, *Bactrocera latifrons*, led to finding a 30 Kb long genomic sequence (Sequence ID: MIMC01001452.1) with an overall 90% MOY protein similarity over a 70 aa long region (BlMoY), suggesting the presence of the whole *MoY* orthologous gene region. A second genomic contig 46 Kb long (Sequence ID: MIMC01001198.1) was also identified coding a shorter BlMOY protein (BlMoY-2), showing 91% MOY protein similarity over a 36 aa long region. A tBLASTn search of 5 available NCBI SRA databases from *B. latifrons* (adult males: SRX1007577; adult females: SRX1007576; embryos: SRX1007578; larvae: SRX1007579; pupae: SRX1007580) failed to find *BlMoY* transcripts. A SRA from embryos showed aa sequence similarity but not identity, suggesting the presence of a transcribed duplicated divergent *BlMoY* gene (SRX1007580; 67% aa similarity over a 31 aa long region). A BLASTp analysis of BlMOY showed a similarity respectively of 95% to BdMOY and of 55% to MOY (over a 60 aa long region).

**BlMOY 70 aa**

```
MGSVWIIIRKHNSRTVILTSSERLILSRKFWNEKNTKPDIEKKEMVLNLCTEFNKLMNNNNKKCLFTRKF

Identities 62/70(89%)
Positives  67/70(95%)
BlMOY  1   MGSVWIIIRKHNSRTVILTSSERLILSRKFWNEKNTKPDIEKKEMVLNLCTEFNKLMNNN  60
           MGSVWIIIRKHNSRTVILTSS+RL+LSR+FWNEKN KPDIE+KE+VLNL TEF KLMNNN
BdMOY  1   MGSVWIIIRKHNSRTVILTSSQRLLLSRRFWNEKNMKPDIEEKEIVLNLSTEFKKLMNNN  60

BlMOY  61  NKKCLFTRKF  70
            NKKCLFTRKF
BdMOY  61  NKKCLFTRKF  70

Identities 18/60(30%)
Positives  33/60(55%)

BlMOY   3       SVWIIIRKHNSRTVILTSSERLILSRKFWNEKNTKPD----IEKKEMVLNLCTEFNKLMN  58
                ++ +I  K+NSRT+ + +S+   +  KFW +          +E+K +V N   E+ KLM+
MOY     11      TISLITIKYNSRTIKVITSKSRGMEPKFWGKMEIAMTENYFVEEKPLVYNFAIEYRKLMS  70
```

**Supplementary Note 11 | *MoY* orthologue in the peach fruit fly *Bactrocera zonata*, which is native of East Asia and present in 20 countries of this area.**

A tBLASTn with BdMOY aa sequence on NCBI SRA databases of another Asian species, *Bactrocera zonata*, led to the finding of a number of SRAs in adult males and pupae (SRX2016848, SRX2016847), but not females neither embryos (SRX2016849, SRX2016846). A tBLASTn with BdMOY protein sequence of a Trinity assembly produced from the downloaded SRA from adult males resulted in the identification of a 0.9 Kb long Trinity transcript encoding for a 70 aa long protein (BzMOY) and showing 93% protein sequence similarity with the probe (TRINITY_DN33806_c0_g3_i1). BLASTp analyses showed that BzMOY (70 aa) is similar to BdMOY (92% aa overall similarity) and to MOY (56% aa similarity over a 60 aa long region) (see below).

**BzMOY 70 aa**

```
MGSVWIIIRKHNSRTVIQTSSERRILSRRIWNEKNTKPDIEKKEMVLNLSTEFKKLMNTNNKKCLFTRKF
Identities 61/70(87%)
Poisitives 65/70(92%)

BzMOY  1       MGSVWIIIRKHNSRTVIQTSSERRILSRRIWNEKNTKPDIEKKEMVLNLSTEFKKLMNTN  60
               MGSVWIIIRKHNSRTVI TSS+R +LSRR WNEKN KPDIE+KE+VLNLSTEFKKLMN N
BdMOY  1       MGSVWIIIRKHNSRTVILTSSQRLLLSRRFWNEKNMKPDIEEKEIVLNLSTEFKKLMNNN  60

BzMOY  61      NKKCLFTRKF  70
               NKKCLFTRKF
BdMOY  61      NKKCLFTRKF  70

17/60(28%)
34/60(56%)

BzMOY  3       SVWIIIRKHNSRTVIQTSSERRILSRRIWNEKNTKPD----IEKKEMVLNLSTEFKKLMN  58
               ++ +I  K+NSRT+   +S+ R +  + W +       +E+K +V N + E++KLM+
MOY     11     TISLITIKYNSRTIKVITSKSRGMEPKFWGKMEIAMTENYFVEEKPLVYNFAIEYRKLMS  70
```

**Supplementary Note 12 | *MoY* orthologue in *Bactrocera correcta*, distributed in Southeast Asia.**

A tBLASTn search of *Bactrocera correcta* pupal and adult male RNA sequence databases (SRA SRX2013590 and SRX2013591) led to finding 2 partially overlapping SRAs, which encode a 35 aa sequence 94% similar to the C-terminus of BtMOY. The 2 SRAs and the truncated BcMOY sequences are reported below.

**BcMOY**
(-) …NQKNTKPDIEKKEMVLNLSTEFKKLMNTNNKKCLF

>gnl|SRA|SRR4020110.18317468.2 FCC6CD9ACXX:3:2114:11885:44523#
CACTTTTTATTATTAGTATTCATTAATTTTTTGAATTCTGTGCTTAAATTCAGTACCATTTCCTTTTTTTCTATATCTGGCTTCGTGTTTTTTT
GATTCC

>gnl|SRA|SRR4020109.22035896.2 FCC6CD9ACXX:3:2303:9390:58342#
GAACAAACACTTTTTATTATTAGTATTCATTAATTTTTTGAATTCTGTGCTTAAATTCAGTACCATTTCCTTTTTTTCTATATCTGGCTTCGTG
TTTTTT

**Supplementary Note 13 | BLASTp and Clustal multiple alignment sequence analysis of MOY orthologues proteins.**

BLASTp analysis of *Ceratitis* MOY protein against 8 other MOY orthologous proteins (Bo, Bd, Bt, Bj, Bl, Bz, Zc and RzMOY) showed a first group of comparable total score of 36-37, in Bo, Rz and Bj species (which are respectively living in the Mediterranean, Australia and North American areas), a second group of comparable total score of 31-28% in other 4 *Bactrocera* and *Zeugodacus* species and a lowest total score (25%) in *Bactrocera latifrons*.

### Supplementary Note 14 | Analysis of the biophysical/structural properties of Ceratitis capitata MoY protein and its orthologs

Biochemical properties of MOY proteins

| Protein | # residues | Mw | Theoretical Isoelectric point |
|---|---|---|---|
| MoY | 70 | 8186 | 9.5 |
| BoMoY | 71 | 8708 | 11.4 |
| BzMoY | 70 | 8467 | 10.8 |
| BlMoY | 67 | 7978 | 9.9 |
| BjMoY | 70 | 8426 | 10.2 |
| BdMoY | 70 | 8468 | 10.5 |
| BtMoY | 70 | 8492 | 10.3 |
| ZcMoY | 60 | 7360 | 10.1 |

Due to the high pI value, all of these proteins are positively charged at neutral pH. Therefore, they are potentially able to interact with negatively charged nucleic acids.

#### *Global/pair-wise alignments and secondary structure prediction by ClustalO alignment*

```
MoY        MDIGNISSKNTISLITIKYNSRTIKVITSKSRGMEPKFWGKMEIAMTENYFVEEKPLVYN 60
ZcMoY      --------MGSVWVLKTKYNSRTI----------TYFFWNAKNTH----LHIETKHIVFN 38
BoMoY      -----MDKMRSVWIIIIKYNSRTVIITTSERRIMPRRVWNAKETK----PHIKKKQMVLN 51
BjMoY      --------MGSVWIIIRKHNSRTVILTSSERLIMSRRFWNEKNMK----PDIEEKEIILN 48
BtMoy      --------MGSVLIIIRKHNSRTVILTSSERLIMSRRFWNEKNMK----PDIEEKEMVLN 48
BdMoY      --------MGSVWIIIRKHNSRTVILTSSQRLLLSRRFWNEKNMK----PDIEEKEIVLN 48
BlMoY      --------MGSVWIIIRKHNSRTVILTSSERLILSRKFWNEKNTK----PDIEKKEMVLN 48
BzMoY      --------MGSVWIIIRKHNSRTVIQTSSERRILSRRIWNEKNTK----PDIEKKEMVLN 48
                        :: ::  *:****:                .*.  :        :: * :: *
Prediction             eeeeeeee   eeeeee                        eee   eeeee

MoY          FAIEYRKLMS--------------- 70
ZcMoY        LTTEFKKLL---NKKCLLTKKFYYK 60
BoMoY        LSTEFKKLK---NKKCLFARKFR-- 71
BjMoY        LSTEFKKLMNNNNTKCLFTRKV--- 70
BtMoy        LSTEFKKLMNNNNKKYLFTRKF--- 70
BdMoY        LSTEFKKLMNNNNKKCLFTRKF--- 70
BlMoY        LCTEFNKLMNNNNKKCLFTRKF--- 70
BzMoY        LSTEFKKLMNTNNKKCLFTRKF--- 70
             :  *:.**
Prediction     ehhhhhhhh             eee
```

Multiple sequence alignments of Moy sequences. Secondary structure prediction was performed using Prediction protocol as implements in PROMALS3D which performs the prediction using multiple sequences. Helices and β-structured regions are denoted with e and h, respectively. As shown in the figure, the analysis of the multiple alignments indicates that there are 11 conserved residues in all sequences. The most conserved region is located in the N-terminal portion of the proteins. Of particular relevance if the hexapeptide KXNSRT that contains two strictly conserved positively charged residues. The lack of sequence similarities with any protein with a known three-dimensional structure makes the determination of MoY putative structural properties difficult. Nevertheless, the reliability of ab-initio secondary structure predictions methods does provide some structural information. PROMALS3D predicts a significant level of secondary structure that for all proteins. Indeed, approximately 60% of the residues of these proteins are embodied in secondary structure elements. This value is in line with that observed for globular compact proteins.

#### *Pair-wise alignments*

Pair-wise sequence identities (%). The numbers in parenthesis represent the similarity (%)

| | Moy | ZcMOY | BoMOY | BzMoy | BlMOY | BjMOY | BdMoY | BtMOY |
|---|---|---|---|---|---|---|---|---|
| MOY | - | 28(57) | 22(41) | 21(41) | 23(41) | 22(44) | 22(44) | 23(44) |
| ZcMOY | | - | 51(70) | 50(70) | 50(69) | 49(68) | 51(69) | 47(65) |
| BoMOY | | | - | 58(64) | 49(57) | 51(60) | 51(62) | 53(60) |
| BzMOY | | | | - | 84(86) | 84(90) | 87(93) | 87(90) |
| BlMOY | | | | | - | 83(89) | 84(91) | 83(87) |
| BjMOY | | | | | | - | 91(97) | 91(94) |
| BdMOY | | | | | | | - | 91(97) |
| BtMOY | | | | | | | | - |

The inspection of the table reporting the pair-wise alignments indicates that the most distant sequence of this ensemble is MoY. BjMOY, BdMOY, and Bt MOY are very similar.
