## Supplementary tables and figures for "*Maleness-on-the-Y* (*MoY*) orchestrates male sex determination in major agricultural fruit fly pests"

| Injection Mix | Injected embryo karyotype | # of injected embryos | # of adults | # of males | # of females | # of intersexes | wt XX females | wt XY males | XY females | XY intersexes |
|---|---|---|---|---|---|---|---|---|---|---|
| #1a: dsRNA targeting *MoY* | XX/XY | 517 | 44 | 11 | 28 | 5 | 25 | 11* | 3 | 5 |
| #1b: dsRNA targeting *MoY* | XX/XY | 200 | 19 | 1 | 18 | 0 | 14 | 1* | 4 | - |
| #1c: dsRNA targeting *MoY* | XX/XY | 500 | 33 | 4 | 27 | 2 | 20 | 4* | 7 | 2 |
| total | XX/XY | 1217 | 96 | 16 | 73 | 7 | 59 | 16 | 14 | 7 |
| buffer only | XX/XY | 110 | 96 | 50 | 25 | 25 | - | - | - | - |

**Supplementary Table 1 | Embryonic *MoY* RNAi induced partial or full feminization of XY individuals.** The *MoY* dsRNA injection experiment #1 reported in Table 1 is the sum of 3 independent injection experiments reported here. In red are indicated numbers of flies showing partial or apparently full sexual transformation. From dsRNA injection experiment #1a: targeting *MoY*, the $G_0$ progeny was composed of apparently normal 25 XX females and 11 XY males, as well as 3 XY females and 5 XY intersexes. The ratio of 25 XX versus 19 XY flies is not significantly deviant with respect to 1:1 ratio (chi-squared test). From dsRNA injection experiment #1b: targeting *MoY*, the ratio of 14 XX versus 5 XY flies is significantly deviant with respect to 1:1 ratio (chi-squared test). In contrast, dsRNA injection experiment #1c showed a ratio of XX versus XY flies close to the expected chi-square value. The male flies marked with *, in female-biased progenies, were assigned to XY karyotype, considering them as *MoY* RNAi escapers.

| ♀ Female | Karyotype | Pupae | Adult flies | males | females |
|---|---|---|---|---|---|
| 26 | XY | 11 | 10 | 1 | 9 |
| 29 | XX | 17 | 10 | | 10 |
| 30 | XX | 0 | 0 | | |
| 31 | XX | 45 | 44 | | 44 |
| 32 | XX | 28 | 24 | | 24 |
| 33 | XX | 26 | 17 | | 17 |
| 34 | XX | 21 | 14 | | 14 |
| 35 | XX | 56 | 54 | | 54 |
| 36 | XX | 6 | 4 | | 4 |
| 37 | XY | 0 | 0 | | |
| 38 | XY | 1 | 0 | | |
| 39 | XY | 0 | 0 | | |
| 40 | XX | 0 | 0 | | |
| 41 | XX | 14 | 11 | | 11 |
| 42 | XX | 0 | 0 | | |
| 43 | XX | 23 | 14 | | 14 |
| 44 | XY | 0 | 0 | | |
| 45 | XX | 0 | 0 | | |
| 46 | XY | 0 | 0 | | |
| 47 | XX | 0 | 0 | | |
| 48 | XX | 0 | 0 | | |
| 49 | XY | 0 | 0 | | |
| 50 | XX | 3 | 3 | | 3 |

| ♀ Female | Karyotype | Pupae | Adult flies | males | females |
|---|---|---|---|---|---|
| 51 | XX | 0 | 0 | | |
| 52 | XX | 2 | 2 | | 2 |
| 53 | XY | 0 | 0 | | |
| 54 | XX | 17 | 15 | | 15 |
| 55 | XX | 0 | 0 | | |
| 56 | XX | 34 | 32 | | 32 |
| 57 | XX | 0 | 0 | | |
| 58 | XX | 0 | 0 | | |
| 59 | XX | 8 | 8 | | 8 |
| 60 | XY | 0 | 0 | | |
| 61 | XX | 0 | 0 | | |
| 62 | XX | 0 | 0 | | |
| 63 | XX | 35 | 32 | | 32 |
| 64 | XY | 0 | 0 | | |
| 65 | XX | 0 | 0 | | |
| 66 | XX | 0 | 0 | | |
| 67 | XY | 0 | 0 | | |
| 68 | XX | 32 | 26 | | 26 |
| 69 | XY | 0 | 0 | | |
| 70 | XY | 0 | 0 | | |
| 71 | XX | 3 | 0 | | |
| 72 | XX | 16 | 14 | | 14 |
| 73 | XX | 0 | 0 | | |

**Supplementary Table 2 | XY $G_0$ females fertility.** Crosses of 46 out of 73 $G_0$ females from dsRNA *MoY* injection (Table 1, #1), each with 3 XX males. The females are numbered referring to the subsequent PCR analysis for karyotyping reported in Extended Data Fig. 4. Only 18 crosses gave $G_1$ progeny. 16 XX females gave no progeny. 17 XX females gave female-only progeny. 11 XY females (in light blue) failed to give any progeny. One XY female (no. 26; in light blue) gave progeny, including 1 male and 9 females, indicating that the Y chromosome can pass through female meiosis and determine male sex. All 46 females were molecularly karyotyped as either XX or XY by PCR on genomic DNA after crossing and 12 XY females here in light blue were identified **(**Extended Data Fig. 4).

| Injected female | Karyotype | Pupae | Adult flies | males | females |
|---|---|---|---|---|---|
| 1 | XX | 70 | 19 | - | 19 |
| 2 | XX | - | - | - | - |
| 3 | XX | 56 | 53 | - | 53 |
| 4 | XX | 31 | 23 | - | 23 |
| 5 | XX | 31 | 23 | - | 23 |
| 6 | XY | 25 | 21 | - | 21 |
| 7 | XX | - | - | - | - |
| 8 | XX | 10 | 8 | - | 8 |
| 9 | XX | - | - | - | - |
| 10 | XX | 2 | - | - | - |
| 11 | XX | - | - | - | - |
| 12 | XX | 63 | 50 | - | 50 |
| 13 | XX | - | - | - | - |
| 14 | XX | 95 | 89 | - | 89 |
| 15 | XX | - | - | - | - |
| 16 | XX | 37 | 30 | - | 30 |
| 17 | XX | 43 | 42 | - | 42 |
| 18 | XX | - | - | - | - |
| 19 | XX | 28 | 27 | - | 27 |
| 20 | XY | - | - | - | - |

**Supplementary Table 3 | $G_0$ XY flies feminized by *Cas9-MoY* embryonic gene disruption can be fertile and transmit *MoY*-defective Y chromosome to $G_1$ progeny.**

20 adult female flies from Cas9-*MoY* injection experiment #3 (Table 1) were individually crossed each with 3 XX males (produced by an *in vivo* RNAi-transgene-mediated masculinization strategy; see materials and methods) and 12 of them gave $G_1$ progeny. PCR karyotyping revealed that 2 out of 20 females were XY (no. 6 and 20). One of the 2 XY females gave a female-only progeny of 21 individuals, as expected in case of *MoY*-defective Y chromosome transmission through the mother. The other XY female (20) died few days after hatching.

| | |
|---|---|
| Total sequenced reads | 179,917,948 |
| Trimmomatic filtered reads | 168,144,620 |
| Ribosomal and mitochondrial depleted reads | 148,886,218 |
| Total assembled bases (bp) | 187,376,939 |
| Trinity assembled transcripts | 213,154 |
| GC content (%) | 37.25 |
| Median transcript length (bp) | 399 |
| Average transcript length (bp) 1253.79 | 879 |
| Transcript N50 (bp) | 1,828 |
| Shortest transcript length (bp) | 201 |
| Longest transcript length (bp) | 17,588 |
| Trinity transcripts > 1Kb | 48,375 |
| Trinity transcripts > 2Kb | 24,380 |

**Supplementary Table 4 | Summary of sequencing and transcriptome assembly statistics.** Illumina short reads sequencing and assembly statistics of the medfly transcript catalogue produced using 4-8 hrs after egg laying embryonic data set and the Trinity *de novo* assembler.

https://www.dropbox.com/s/n8fru9p13vuz5it/Supplementary%20Table%205.xlsx?dl=0

**Supplementary Table 5 | RNA-seq expression values of 4-8h medfly embryonic transcriptome.** Transcript-level quantifications of the 4-8h medfly embryonic transcriptome for each sample (XYXX and XX-only embryos) were done against the assembled transcriptome using Kallisto software[36].

https://www.dropbox.com/s/wp4wyfk486jcgp2/Supplementary%20Table%206.xlsx?dl=0

**Supplementary Table 6 | Results of the differentially expression analysis.** Differential gene expression analysis was performed using edgeR software[37], with cut-off values of FDR < 0.05 and logFC > 0.

| ID | length | FAM18 male_counts | FAM18 female_counts | ISPRA male_counts | ISPRA female_counts |
|---|---|---|---|---|---|
| Cctra-2 (single copy, autosomal gene) | 1113 | 31 | 25 | 40 | 48 |
| pY114 (Y-specific repetititve element) | 1405 | 160 | 0 | 232 | 0 |
| pm11 (Y-specific repetititve element) | 2733 | 148*** | 0 | 317*** | 0 |
| 5Kb (Y-specific repetititve element) | 5642 | 0*** | 0 | 2344*** | 6 |
| pM21 (Y-specific repetititve element) | 1380 | 189*** | 0 | 327*** | 0 |
| Total reads | | 242122621 | 246407224 | 322938668 | 275803598 |

**Supplementary Table 7 | Summary of results of the mapping analysis performed to verify the reduced complexity of the medfly Y chromosome in FAM18 strain versus ISPRA strain.** We compared the read counts obtained by Bowtie mapping of the FAM18 genomic male and female reads on four Y-specific repetitive elements with the reads counts obtained by Bowtie mapping results obtained with publicly available male and female genomic Illumina reads (SRR847687 and SRR847380 for males, SRR847688 and SRR847689 for females) of ISPRA medfly strain, produced at Baylor College of Medicine. We utilized as mapping control the single-copy gene *Cctra-2*. *** $p < 0.001$, Fisher Exact Test.

https://www.dropbox.com/s/sgxpulcpyien1e8/Supplementary%20Table%208.xlsx?dl=0

**Supplementary Table 8 | Summary of transcriptome assembly statistics and SRA accession numbers of Tephritidae species.** Illumina RNA-seq data for 14 *Bactrocera* species were downloaded from NCBI SRA archive and *de novo* assemblies were produced using Trinity assembler[32,33] with default parameters.

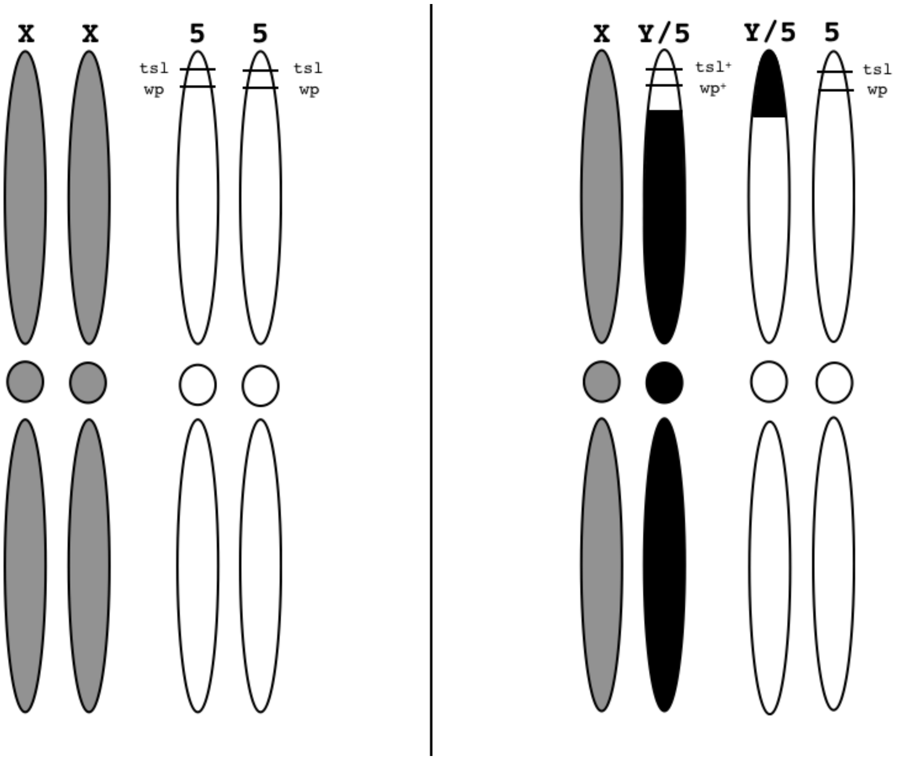


**Supplementary Fig. 1 |** In the TSL strain (FAO-IAEA, Seibersdorf, Austria), females carry the autosomal *tsl* (temperature sensitive) and *wp* (white pupae) mutant recessive alleles in homozygosity, while males carry a reciprocal autosome–Y translocation, with the segment on the Y having the corresponding wild type alleles (*tsl*+ and brown pupae *wp*+).

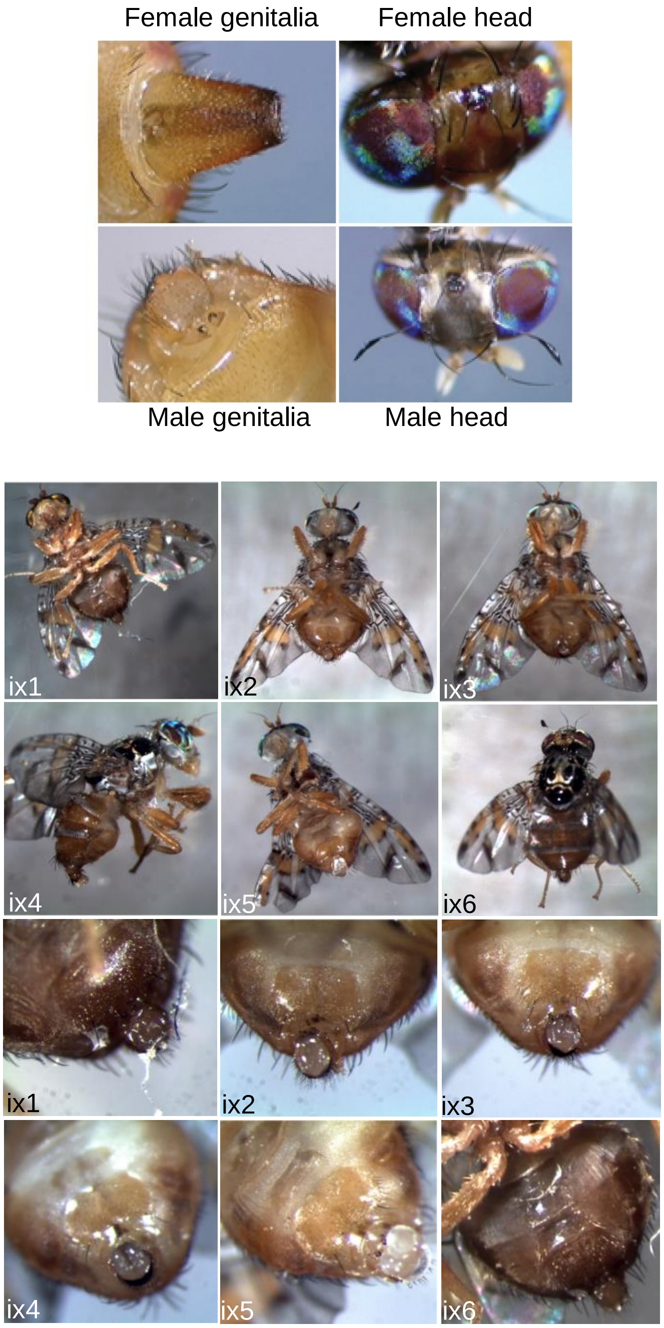


**Supplementary Fig. 2 | Partial phenotypic masculinization of XX individuals induced by *MoY* 5 Kb genomic region.** Upper panel: wild type male and female head and genitalia. Lower panel: 6 XX intersexes from *MoY* DNA embryos injection set #4 (Table 1, set #4; Extended Data Table 3, set #4b; molecular karyotyping not shown) showed male external genitalia, but no male-specific setae on their head (ix1, ix3, ix4 and ix5) or only one setae (ix2 and ix6). We concluded that the intersexes are partially masculinized individuals.

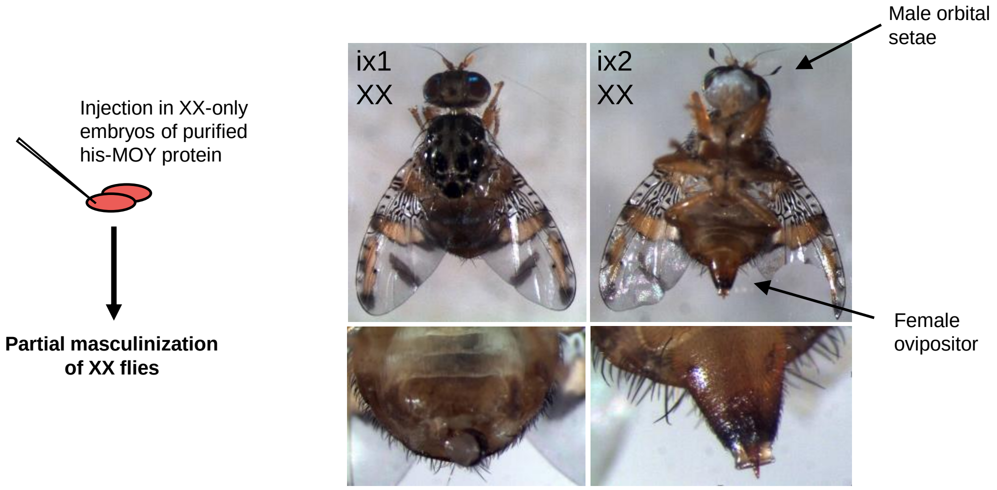


**Supplementary Fig. 3 | Embryos injections of** MOY purified HIS-tagged protein induces partial masculinization in anterior or posterior fly body regions.
