## Extended data tables and figures for "*Maleness-on-the-Y* (*MoY*) orchestrates male sex determination in major agricultural fruit fly pests"

| N° | DE+CQ transcripts 4-8h | Medfly | Baylor Genome | Presence in Fam18 Canu | male-specific in 4-8 h Ben embryos | CQ transcripts 0-48h+males | Blastx Ceratitis | Blastx Drosophila | male specific in B. oleae embryos | mapdXY.CQ | mapdXX.CQ | CQ |
|---|---|---|---|---|---|---|---|---|---|---|---|---|
| 1a | TRINITY_DN40516_c0_g1_i6 | No | (shorter highly related sequences) | multiple | Yes | | no similarity | no similarity | no similarity | 11118 | 0 | 0 |
| 1b | TRINITY_DN40516_c0_g2_i3 | No | (shorter highly related sequences) | multiple | Yes | Lyra | putative gustatory receptor 59f XP_004526066.1 (E value 2.9) | none | no similarity | 10916 | 0 | 0 |
| 1c | TRINITY_DN40516_c0_g2_i2 | No | (shorter highly related sequences) | multiple | Yes | Lyra | putative gustatory receptor 59f XP_004526066.1 (E value 1.7) | zpg | no similarity | 10794 | 0 | 0 |
| 1d | TRINITY_DN40516_c0_g1_i2 | No | (shorter highly related sequences) | multiple | Yes | Lyra | no similarity | no similarity | no similarity | 10759 | 0 | 0 |
| 1e | TRINITY_DN40516_c0_g2_i1 | No | (shorter highly related sequences) | multiple | Yes | | putative gustatory receptor 59f XP_004526066.1 (E value 2.2) | mabiki [Drosophila melanogaster] (E value 7.5) | yes, short and weak | 11935 | 0 | 0 |
| 2 | TRINITY_DN38563_c5_g1_i1 | No | (shorter highly related sequences) | none | Yes | | branchpoint-bridging protein (E value 6e-39) | quaking related 58E-2, isoform A (E value 1e-30) | no similarity | 79 | 0 | 0 |
| 3a | TRINITY_DN40292_c0_g1_i10 | No | (shorter highly related sequences) | multiple | Yes | | no similarity | no similarity | no similarity | 5813 | 0 | 0 |
| 1f | TRINITY_DN40516_c0_g1_i5 | No | (shorter highly related sequences) | none | Yes | | no similarity | no similarity | no similarity | 10249 | 0 | 0 |
| 3b | TRINITY_DN40292_c0_g3_i1 | No | No | single | Yes | Corvus | none | Lace isoform E (E value 1.7) | yes, short | 68 | 0 | 0 |
| 4a | TRINITY_DN40142_c1_g4_i1 | No | (highly related paralogous sequences) | multiple | Yes | | histone H2B (E value 6e -43) | histone H2B (E value 2e-42) | no-male specific | 872 | 0 | 0 |
| 5 | TRINITY_DN33215_c0_g2_i1 | No | (highly related paralogous sequences) | none | No | | uncharacterized protein LOC105665391 (E value 5.6) | none | yes, short | 3982 | 0 | 0 |
| 1g | TRINITY_DN40516_c0_g1_i7 | No | (shorter highly related sequences) | multiple | Yes | | no similarity | no similarity | none | 8580 | 0 | 0 |
| 6 | TRINITY_DN40470_c17_g1_i5 | No | (highly related paralogous sequences) | none | Yes | | cytosol aminopeptidase (E value 2e-23) | Sperm-Leucylaminopeptidase 5 (E value 3e-13) | yes | 3182 | 0 | 0 |
| 4b | TRINITY_DN40142_c1_g1_i5 | No | (highly related paralogous sequences) | multiple | No | | no similarity | no similarity | yes | 274 | 0 | 0 |
| 4c | TRINITY_DN40142_c1_g4_i2 | No | (shorter highly related sequences) | multiple | Yes | | histone H2B (E value 9e-30) | histone H2B (E value 1e-29) | no-male specific | 336 | 0 | 0 |
| 7 | TRINITY_DN38104_c3_g2_i2 | Yes | No | none | No | | no similarity | no similarity | none | 6253 | 0 | 0 |
| 8 | TRINITY_DN40402_c3_g5_i1 | No | (highly related paralogous sequences) | multiple | Yes | | no similarity | no similarity | none | 16013 | 0 | 0 |
| 9 | TRINITY_DN36540_c0_g1_i1 | Yes | No | none | No | | no similarity | no similarity | none | 11903 | 0 | 0 |
| 10 | TRINITY_DN37671_c9_g1_i1 | Yes | No | none | Yes | | F-Box/SPRY domain-containing protein (E vaule 9.4) | no similarity | none | 11015 | 0 | 0 |

**Extended Data Table 1 | List of 19 embryonic (4-8 h) transcripts corresponding to 10 putative Y-linked genes, filtered by DE and CQ analyses.** BLASTn analyses showed that most of the 19 transcripts are missing in the currently available medfly genome (NCBI), which however contain paralogous sequences. Only 3 transcripts have 100% corresponding genomic sequences in this assembly. In contrast, BLASTn analyses on the male medfly Canu *Fam18* genome showed that 12 transcripts have corresponding identical sequences, with most present in different contigs suggesting multiple copies. The presence of corresponding transcripts only in the mixed XX/XY but not in the XX-only transcriptome further supported male-specificity for 15 of them. 4 out of 19 transcripts corresponded to 2 previously selected putative Y-linked male-specific genes (*lyra* and *corvus*). BLASTx analysis on protein databases of *C. capitata* and *D. melanogaster* showed some similarity mostly to short stretch of peptidases, transcriptional factors, receptors and histone proteins. The number of *Fam18* male (mapdXY.CQ) and female (mapdXX.CQ) mapped reads and the chromosome quotient value (CQ), calculated as mapdXX.CQ/mapdXY.CQ, are reported.

| Injection Mix | Karyotypes of injected embryos | Injected embryos | Adults | XY males | XX females | XY females | XX males | XY intersexes | XX intersexes |
|---|---|---|---|---|---|---|---|---|---|
| #1: dsRNA *MoY* | XX/XY | 1217 | 96 | 16* | 59 | **14** | 0 | **7** | 0 |
| #2: dsRNA *MoY* | X X; *wp/wp*<br>X Y-*wp*$^{+}$; *wp,* A-Y | 260 | 10 | 2* **(**XY-*wp*$^{+}$) | **7** | **1** **(**XY-*wp*$^{+}$) | 0 | 0 | 0 |
| #3: CRISPR/Cas9 vs *MoY* | XX/XY | 250 | 32 | 7* | 18 | **2** | 0 | **5** | 0 |
| #4: linear 5 Kb *MoY* DNA | XX/XY | 310 | 28 | 16 | 3° | 0 | **3** | 0 | **6** |
| #5: *MoY* 5 Kb plasmid | XX/XY | 190 | 16 | 8 | 5° | 0 | **1** | 0 | **2** |
| #6: his-MOY protein | XX | 428 | 31 | - | 25 | - | 0 | - | **6** |
| #7: dsRNA *BoMoY* | XX/XY | 550 | 24 | 6 | 10 | **3** | 0 | **5** | 0 |
| #8: dsRNA *BdMoY* | XX/XY | 540 | 41 | 16 | 17 | **4** | 0 | **4** | 0 |

**Extended Data Table 2 | *MoY* is necessary (#1, #2 and #5) and sufficient (3#, 4# and 6#) for male sex determination and functionally conserved in *Bactrocera* species (#7-#8).** Medfly embryos injections at 0-1 h AEL of *MoY* dsRNA (#1-#2), DNA (#3, #4), Cas9 RNP (#5), and protein (#6). Embryos injections of *MoY* orthologues dsRNA in *Bactrocera oleae* and *B. dorsalis* (8#, 9#). In red are indicated numbers of flies showing partial or apparently full sexual transformations. In injection set 2#, a Y-marked brown pupae strain, carrying a white pupae recessive mutation on an autosome was used (reciprocal autosome-Y chromosome translocation). The male flies marked with * in strongly female-biased progenies (#1, #2 and #5) were assigned to XY karyotype without molecular analyses, considering them as *MoY* RNAi or Cas9 escapers. Similarly, the female flies marked with ° in male-biased progenies (#3 and #4) were assigned to XX karyotype without molecular analyses. Injection set 1# reports data from 3 biological replicates reported in Extended Data Table 2 .

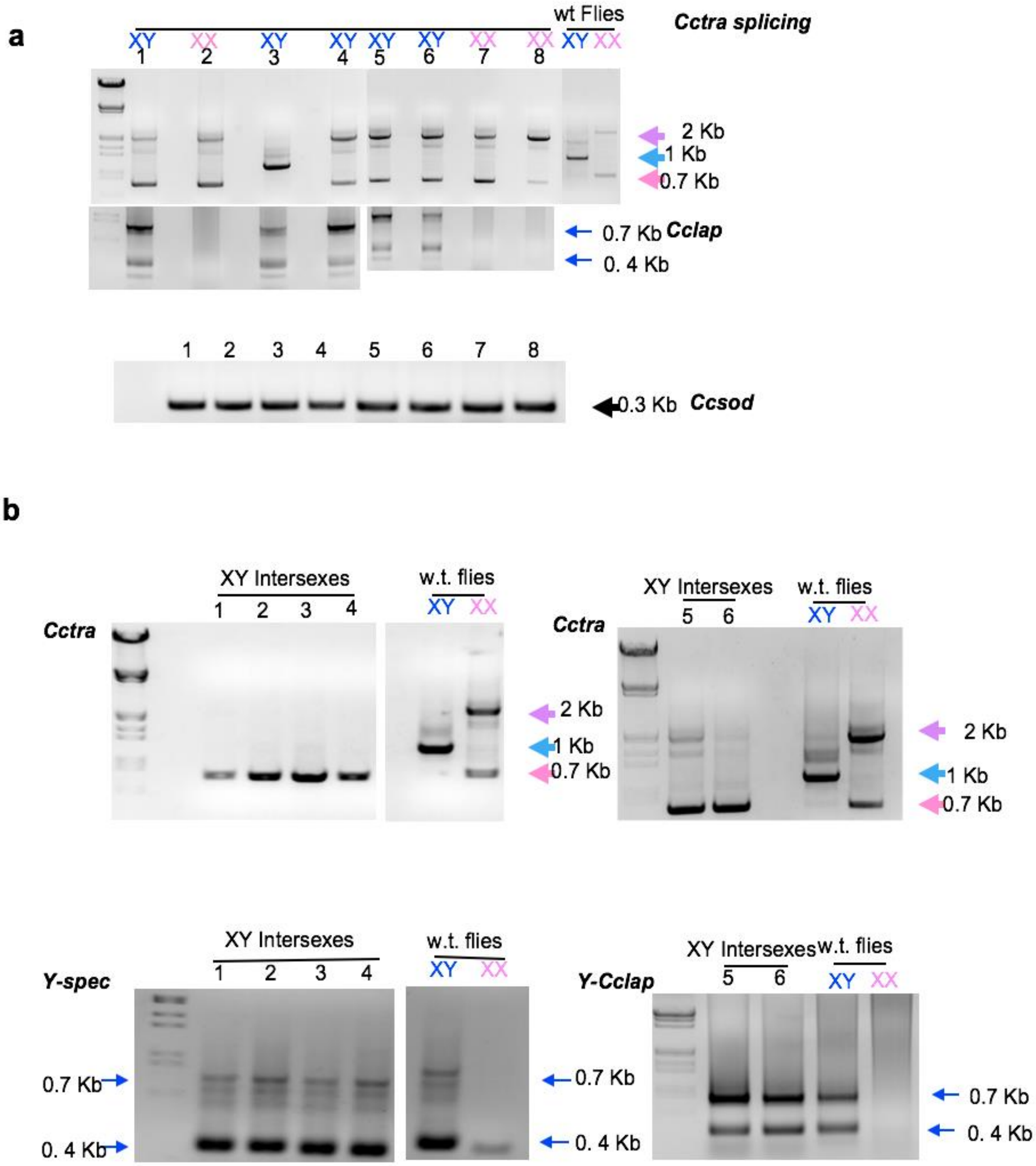


**Extended Data Fig. 1 | Transient embryonic silencing of *MoY* leads to *Cctra* female-specific splicing in XY larvae. a,** RT-PCR analyses of *Cctra* in 8 larvae hatched from *MoY* dsRNA-injected embryos showing *Cctra* female-specific transcripts (2 Kb and 0.7 Kb cDNA bands) in 4 out of 5 XY individuals (lanes 1, 4, 5 and 6). No effect was observed on female-specific *Cctra* splicing pattern in 3 XX larvae (lanes 2, 7 and 8) and on male-specific splicing pattern of 1 XY (lane 3) larvae. Sex-specific *Cctra* transcripts were amplified from adult flies as a reference for gel migration of the corresponding cDNA bands (female-specific 2.1 Kb and 0.7 Kb; male-specific 1.1 Kb). Individual larvae were molecularly karyotyped using a Y-derived transcript from *Cclap* pseudogene (Salvemini et al., 2011). *Ccsod* transcripts were used as a positive control and as negative control for genomic DNA contamination. **b,** RT-PCR analyses of *Cctra* carried out in 6 out of 7 adult intersex XY flies showed either female-specific transcripts (1-4) or a mix of male-specific and female specific ones (5-6). Molecular karyotyping of XX/XY individuals by RT-PCR on Y-specific transcribed sequences (transcribed repetitive Y-linked sequence pY114–related by YF/YR primers; lanes 1-4; Anleitner and Haymer, 1992) and Y-linked *Cclap* (lanes 5-6) confirmed the expected XY karyotype of the intersexes, indicating partial feminization. Sex-specific *Cctra* and Y-specific *Cclap* transcripts were amplified from adult flies as a reference for gel migration of the corresponding cDNA bands.

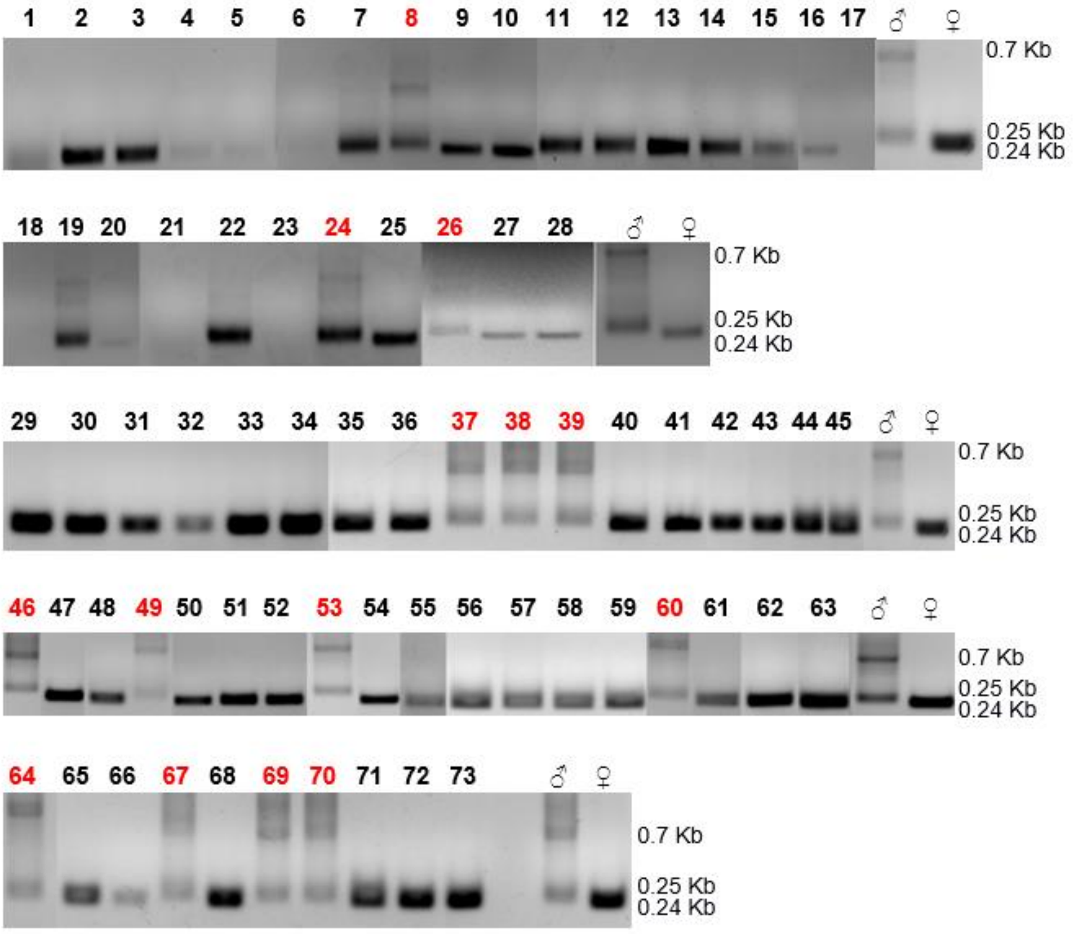


**Extended Data Fig. 2 | Molecular karyotyping of 73 G0 females obtained from embryonic *MoY* RNAi.** Sexing by PCR karyotyping was performed on genomic DNA from a small wing fragment dissected from each of 73 females from injection set #1 (Table 1). The presence of the Y chromosome (as 0.7 Kb and 0.250 Kb bands) was detected in 14 out of 73 adult females (in red) using *CcYF/CcYR* primers[31]. In the remaining 59 adult females, a slightly smaller band was detected indicating the absence of the Y (0.24 Kb) [31].

a

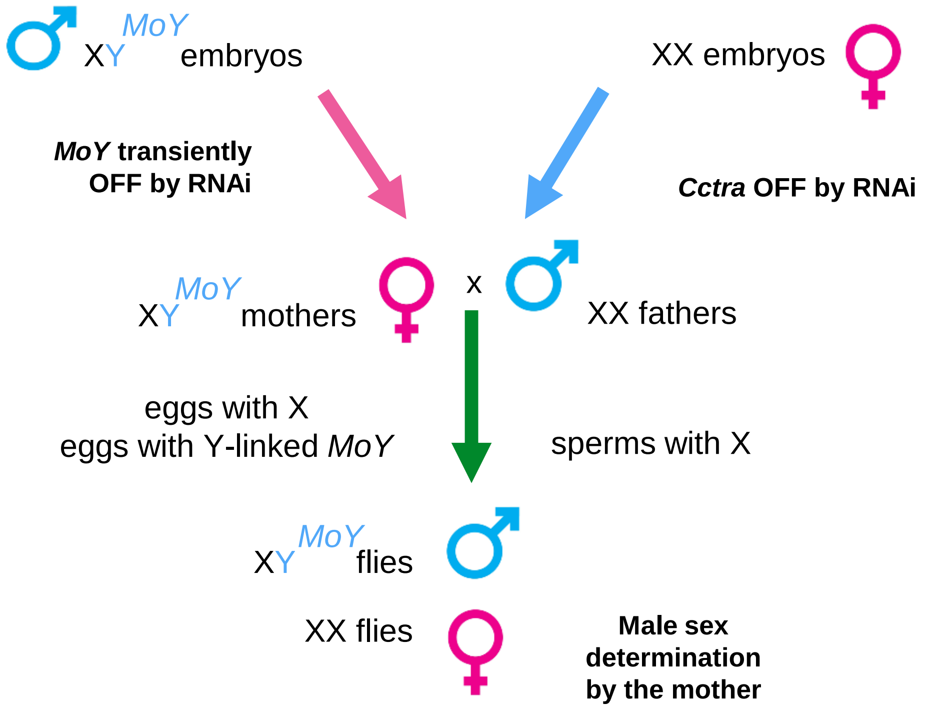


b

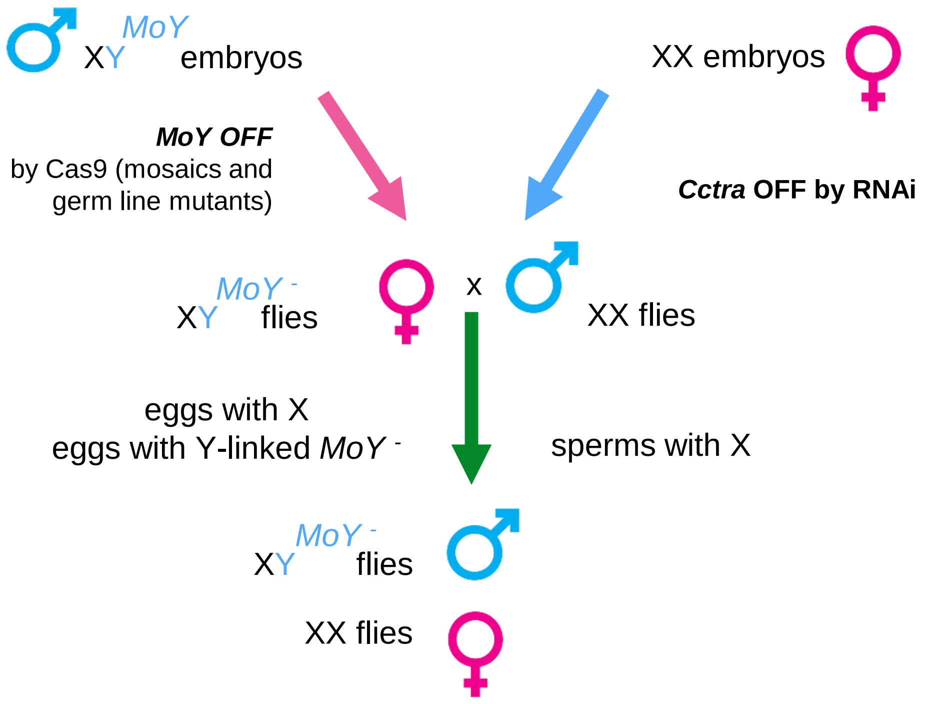


**Extended Data Fig. 3 | In *Ceratitis capitata*, artificial transient female heterogamety (XY) and male homogamety (XX) are compatible with fertility and male sex determination. a,** XY mothers were obtained by transient *MoY* embryonic RNAi. XX fathers were obtained by transient *Cctra* RNAi (see Methods). *MoY* gene can be transmitted by the mother and determines the male sex in the progeny. **b,** Maternal transmission of a Y chromosome carrying a *MoY* Cas9-induced null allele.

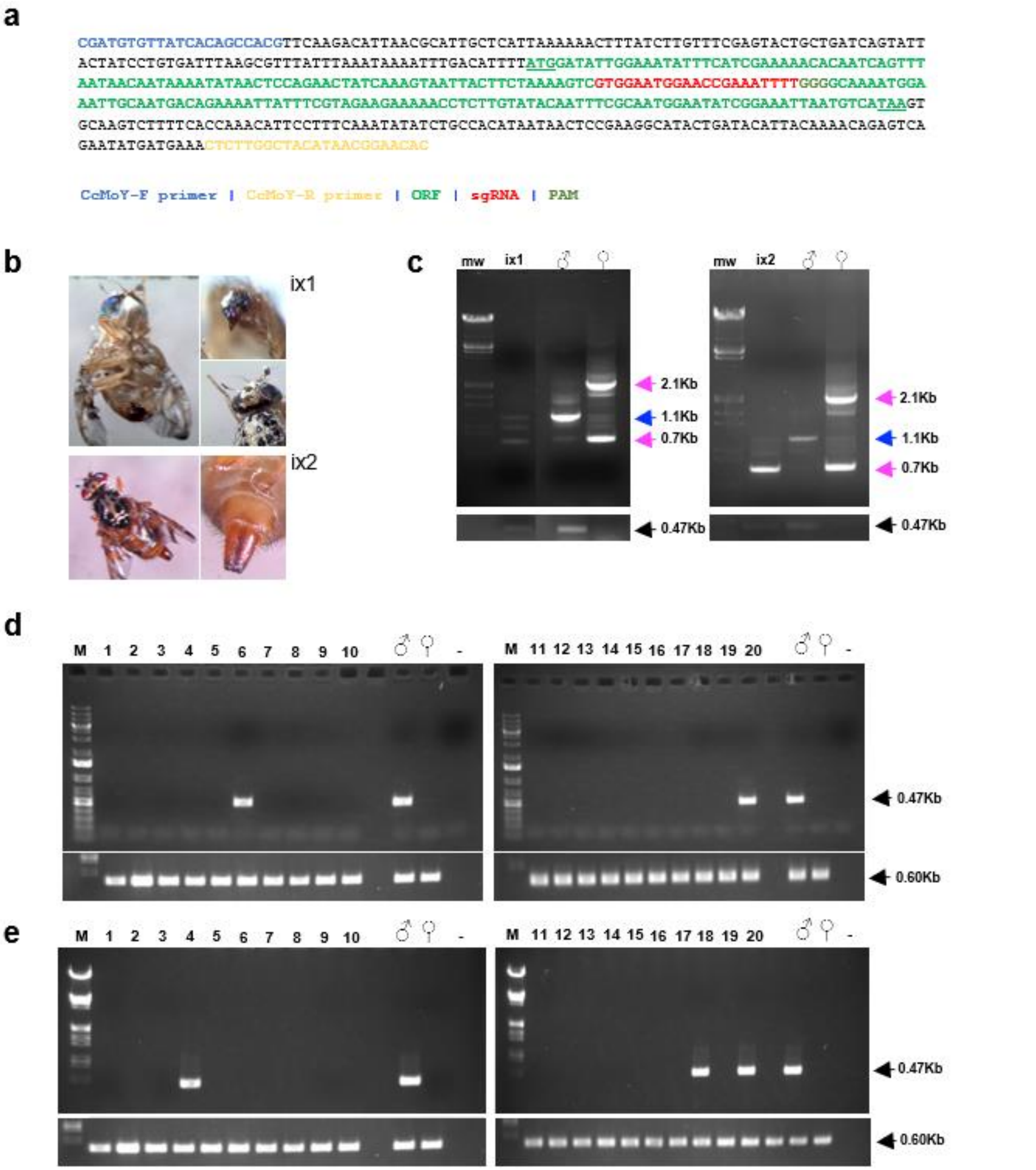


**Extended Data Figure 4 | CRISPR-induced partial feminization of XY individuals. a**, *MoY* coding region sequence, Cas9 target site, and primers used for PCR. ATG and STOP codon are underlined. **b**, 2 adult intersexes developed after embryonic CRISPR-Cas9 injections: ix1 shows male-head and malformed ovipositor; ix2 shows male-head and female ovipositor. **c**, RT-PCR of *Cctra* and of *MoY* (0.47 Kb) for karyotyping: both intersexes shown in figure b are XY and express a mix of male and female *Cctra* transcripts. **d**, Molecular karyotype analysis on the 20 $G_0$ female flies developed from Cas9-gRNA embryonic injections. The sexing by PCR karyotyping was performed on genomic DNA from adult females injected with the ribonucleoprotein Cas9/*MoY*-gRNA complex, using primers CcMoY-F and CcMoY-R. Two females (numbers 6 and 20) were found to be positive for *MoY*, showing a complete feminization due to the injection of the ribonucleoprotein Cas9/*MoY*-gRNA complex, causing a knock-out of the *MoY* gene. **e**, Molecular karyotype analysis on 21 $G_1$ female flies born from female 6 from Cas9-*MoY* injected embryos (Table 1 - Figure 3). Sexing by PCR karyotyping was performed on genomic DNA from the 21 females born from female 6 (see Table 1), using primers CcMoY-F and CcMoY-R. Three females (numbers 4, 18 and 20) were found to be positive to *MoY*, thus inherited the knock-out *MoY* gene from the mother. *CcSOD* PCR on genomic DNA, used as positive control, shows that the genomic DNA of all samples was amplifiable in both d and e panels.

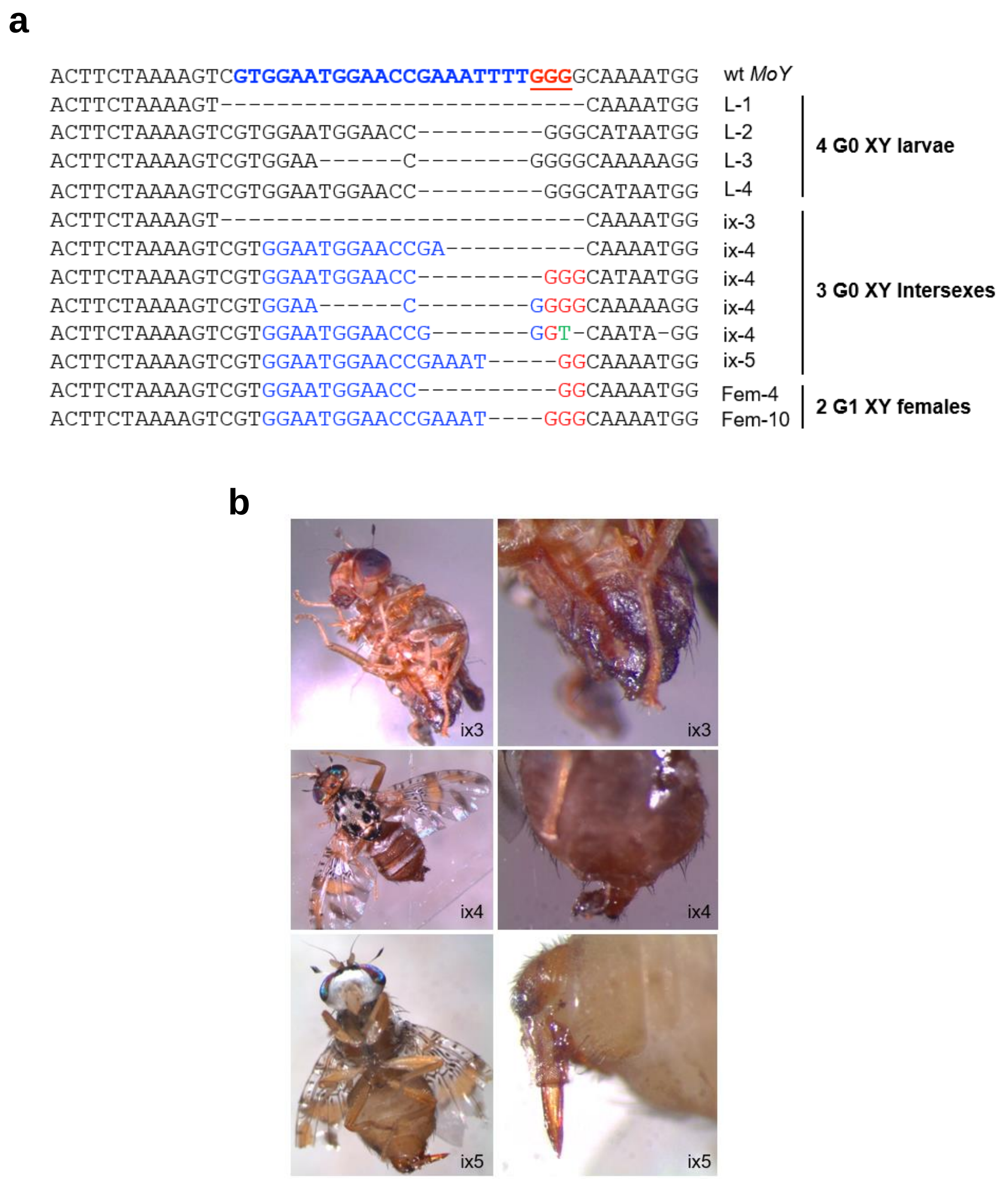


**Extended Data Figure 5 | CRISPR-Cas9–induced disruption of *MoY* causes complete male-to-female transformation of XY individuals. a,** Injection of Cas9 ribonucleoparticles targeting the putative coding region of *MoY* (see Fig. 1d; Table 1, #5) induced indels in proximity to the PAM site, as detected in DNA from a pool of $G_0$ larvae and from 3 adult $G_0$ intersexes (see also Extended Data Fig. 12). One intersex is shown with feminization of the head region (no male-specific setae). Two $G_1$ XY females (female 4 and 20, (born from a full feminized $G_0$ XY *MoY*-CRISPR*ant* mother and a XX "special" father), showed deletions in *MoY* of respectively 10 bp and 4 bp, causing frameshift mutations (Table 1, #3). **b**, Cas9-induced intersexual phenotypes. Additional three intersexes showed the presence of male bristles on the head (left column) and deformed genitals similar to ovipositors (right column). Indels mutations in the targeted *MoY* region in 3 adult intersexes (ix3, ix4 and ix5) are shown in Fig. 4

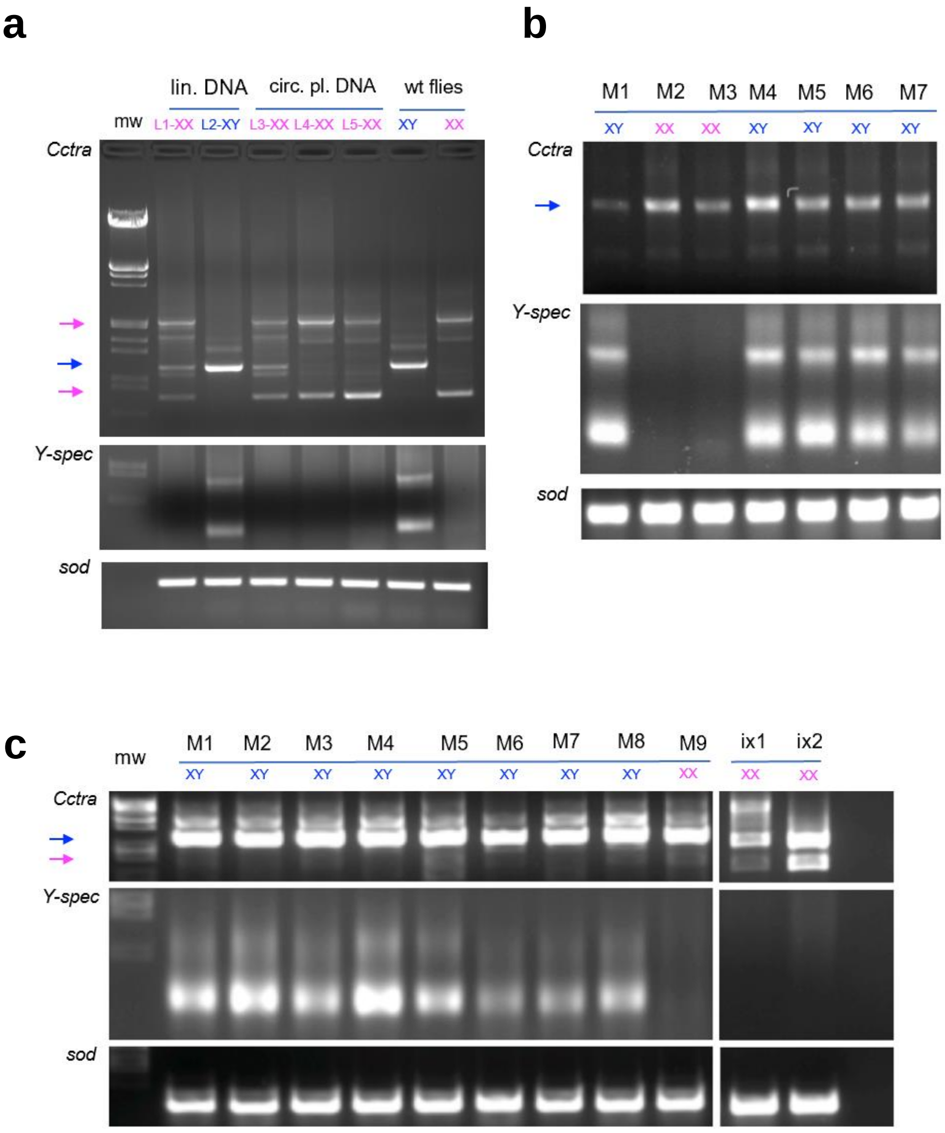


**Extended Data Fig. 6 | Molecular masculinization of XX larvae following embryos injection with *MoY* genomic DNA.** **a**, **b**, **c**, RT-PCR of *Cctra*, the Y-linked *Cclap* (b) and *Ccsod* as positive control (c) on individuals from injected embryos (Extended Data Table 3, injection set 3# and 4#)(mw: molecular marker). **a,** L1 and L3 larvae, respectively from injected embryos with linear (Lin.) or plasmidic (circ. = circular)) *MoY* DNA showed XX karyotype (lack of Y-specific *Cclap* cDNA bands) and a male-specific *Cctra* band (blue arrow; 1.1 Kb), in addition to the female specific ones (pink arrows, 2 Kb and 0.7 Kb). No effect is observed in XY larvae. L4 and L5 XX larvae are escapers of the masculinization induced by the *MoY* plasmid, possibly because of quantitative variability in the manual embryos injections. Some minor *Cctra* different bands visible in distinct lanes are likely due to variable amplification of intermediates of *Cctra* splicing. **b**, Two out of 7 adult males (Extended Data Table 3, *MoY* DNA linear fragment embryos injections set 3#,), showed XX karyotype but male-specific *Cctra* splicing product (blue arrow; 1.1 Kb), indicating apparently full molecular masculinization. Molecular karyotyping (Y-spec) and positive control (sod) are also shown. Similar molecular analysis and karyotyping were performed on the remaining 12 males from the same injection set 3# (data not shown). **c**, One of 9 adult males from *MoY* DNA plasmid embryos injection set 4# (Extended Data Table 3) showed XX karyotype and male-specific *Cctra* splicing, suggesting again full molecular masculinization. Two phenotypic XX intersexes showed a mix of male- and female-specific *Cctra* products.

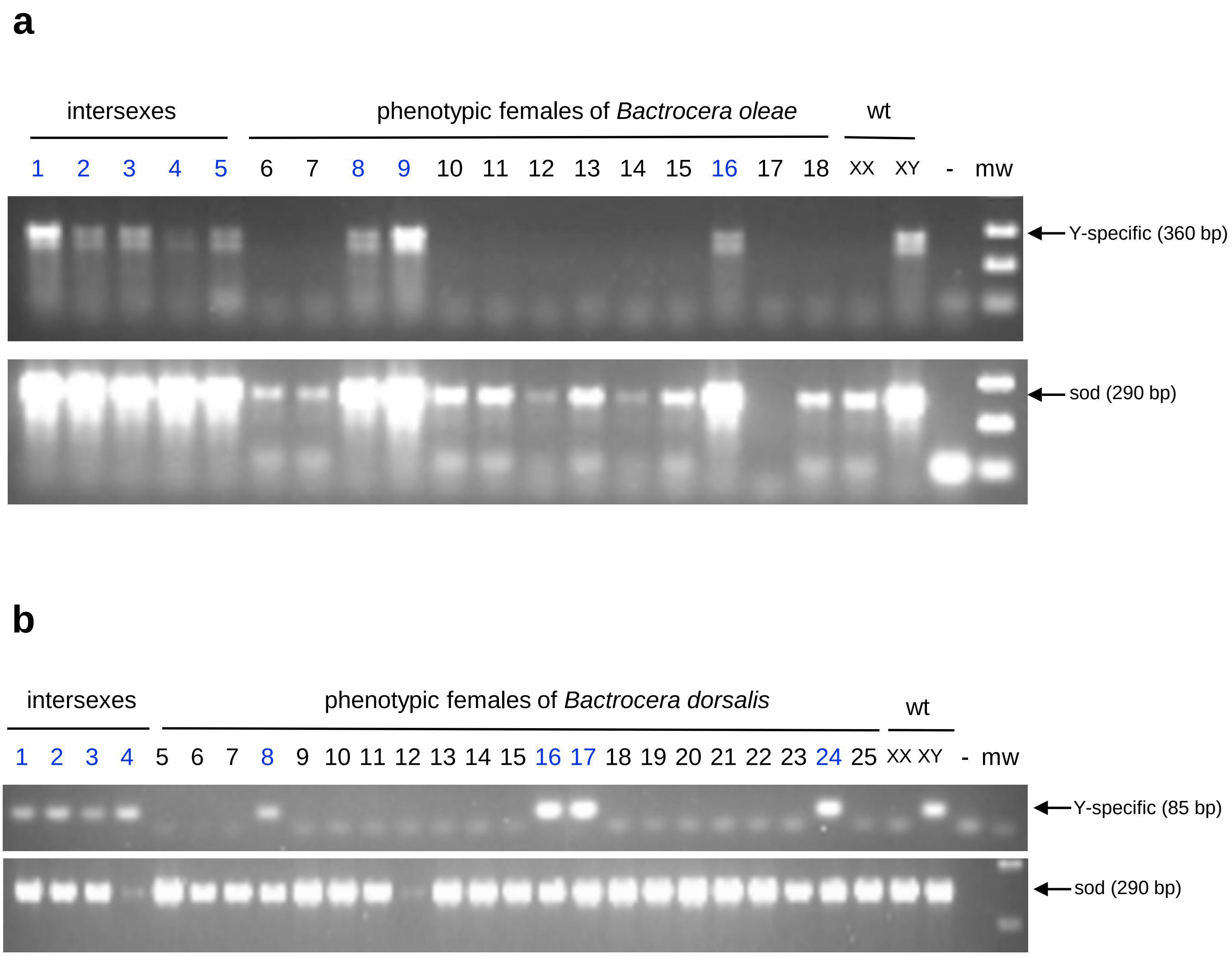


**Extended Data Figure 7 | Molecular karyotyping of intersexes and G0 females from *MoY* embryonic RNAi in *B. oleae* (a) and *B. dorsalis* (b).** Adult flies from injections sets 7# and 8# were molecularly karyotyped by PCR with Y-specific primers (see mat. and meth.). **a,** 5 *B. oleae* intersexes and 3 females were found to be XY. XX and XY are positive controls (the female and male Bo adults, respectively).**b,** 4 *B. dorsalis* intersexes and 4 females were found to be XY. XX and XY are positive controls (the female and male Bd adults, respectively).
